# RNA-seq meta-analysis and machine learning identify stress-responsive genes and improve genomic prediction in common bean (*Phaseolus vulgaris* L.) with cross-species application in cowpea (*Vigna unguiculata* L.)

**DOI:** 10.64898/2026.08.08.743654

**Authors:** Dotun Olaoye, Laide Rasaki, Olamide Adesina, Babatunde Kareem, Shyam Kandel, Waltram Ravelombola, Yufeng Yang, Ainong Shi

**Affiliations:** Department of Horticulture, University of Arkansas, Fayetteville, AR, USA; Department of Crop and Soil Sciences, North Carolina State University, Raleigh, NC, USA; Department of Plant Pathology, Kansas State University, Manhattan, KS, USA; Department of Plant Pathology and Plant-Microbe Biology, Cornell University, Ithaca, NY, USA; USDA-ARS, Edward T. Schafer Agricultural Research Center, Sugarbeet Research Unit, Fargo, ND 58102, U.S.A; Genetics Area Program, University of Missouri, Columbia, MO, USA

**Keywords:** Abiotic stress, Biotic stress, Meta-analysis, Machine learning, Genomic selection

## Abstract

Common bean (*Phaseolus vulgaris* L.) is exposed to a broad spectrum of abiotic and biotic stresses that impose severe constraints on productivity, yet the molecular basis of stress tolerance remains poorly resolved, with independent studies yielding inconsistent and incomplete conclusions. To establish a comprehensive picture of the common bean stress transcriptome, we conducted a systematic meta-analysis of publicly available RNA-sequencing datasets spanning abiotic and biotic stress conditions across leaf and root tissues. Integrating statistical meta-analysis with machine-learning approaches, we identified stress-responsive gene sets whose robustness was verified through rigorous statistical approaches including independent dataset validation. Beyond confirming established stress-responsive genes, the machine-learning framework uncovered candidates overlooked by standard significance thresholds in individual studies yet carrying consistent transcriptional signals across studies. Co-expression and protein-protein network analyses further resolved these candidates into functionally coherent modules linked to specific stress-response programs. Notably, ethylene-responsive transcription factors were identified as hub genes in three of four stress-tissue groups, with NAC domain transcription factors emerging as additional hub genes in biotic stress contexts. Importantly, the biological significance of the identified gene sets was validated genomically: marker panels targeting consensus meta-analysis-derived and machine-learning-discovered gene regions improved genomic prediction accuracy for disease resistance traits in common bean and abiotic stress tolerance traits in cowpea relative to a baseline model with equivalent-sized random marker sets. Overall, these findings revealed conserved stress transcriptome signatures in common bean and provided a cross-species, evidence-based framework for prioritizing candidate genes and constructing biologically informed genomic selection tools to advance stress-resilient legume breeding.

## Introduction

Common bean (*Phaseolus vulgaris* L.) is one of the most widely cultivated pulse crops globally, valued primarily as a source of dietary protein for smallholder farming communities in sub-Saharan Africa, Latin America, and South Asia (Akibode and Maredia, 2012; Beebe et al., 2013; Camara et al., 2013). Annual production losses attributable to drought can be up to 80% depending on growth stage and stress duration, with critical impacts directly halting the flowering and pod-filling processes (Polania et al., 2016). Furthermore, severe outbreaks of plant pathogens can result in total field loss when environmental conditions favor high disease severity on a susceptible cultivar (Smith et al., 2019). These losses are projected to increase as climate change intensifies the frequency and severity of abiotic and biotic stress events. Thus, the identification of molecular targets for stress tolerance improvement remains a priority for common bean breeding programs.

Plants respond to abiotic and biotic stresses through coordinated molecular mechanisms involving stress signal perception, hormonal signaling, transcriptional reprogramming, and metabolic adjustment. Abiotic stresses including drought, heat, cold, and salinity disrupt key physiological processes such as stomatal closure, reduction in carbon assimilation, distorted assimilate partitioning, and accumulation of reactive oxygen species (ROS), initiating a molecular cascade that spans from initial stress perception through progressive cellular damage (Wu et al., 2014). Biotic stresses imposed by pathogens like fungi and bacteria activates pattern-triggered and effector-triggered immunity, inducing the expression of pathogenesis-related genes. The plant hormones abscisic acid (ABA), jasmonic acid (JA), salicylic acid (SA), and ethylene act as central coordinators of these responses, with their crosstalk determining the balance between growth and defense under stress (Pieterse et al., 2009; Verma eta al., 2016).

Stress responses in common bean are not uniform across tissues or conditions. Root and leaf tissues experience distinct stress inputs and mount correspondingly different transcriptional responses: roots encounter soil-borne pathogens, osmotic imbalance, and ion toxicity directly, while leaves are the primary site of photosynthesis and are more directly exposed to atmospheric drought, heat and pathogens. Transcriptomic analyses in related plant species show that root and shoot stress responses differ substantially in both the identity and magnitude of differentially expressed genes, with relatively few elements shared between tissues (Pereira et al., 2020). Similarly, abiotic and biotic stress responses, while overlapping in ROS signaling and cell wall modification, diverge in hormone signaling and effector gene expression (Leitão et al., 2021). Thus, studies restricted to a single tissue or stress type cannot capture the full landscape of stress-responsive gene regulation in common bean.

Despite the specificity of individual stress responses, evidence from multiple plant species indicates that a shared molecular core is activated across distinct stress types. Stress priming, which is a brief exposure to a mild stressor that subsequently confers tolerance to an unrelated stress, demonstrates cross-tolerance between drought and heat, cold and salinity, and abiotic and biotic stressors, implying the activation of common molecular defense mechanisms (Balmer et al. 2015). Considerable effort has been directed at understanding stress responses in common bean through transcriptome profiling across drought, pathogen infection, and other conditions in leaf and root tissues (Torres et al., 2006; Wu et al., 2014, 2017; Gregorio Jorge et al., 2020; Pereira et al., 2020), yet findings from independent studies are difficult to integrate because they differ in genotype, tissue, growth stage, and treatment protocol. Whether a core of genes responds consistently across abiotic and biotic stress conditions and tissue types in common bean remains unresolved, and establishing it is a prerequisite for identifying the most robust targets for genetic improvement.

Standard differential expression meta-analysis recovers only genes that meet significance thresholds consistently across all contributing studies. Machine learning (ML) classifiers can extend this gene set by identifying genes with expression patterns characteristic of stress response even when they fall short of most-common statistical significance thresholds. Support vector machine (SVM) model trained on pre-labeled gene sets have been applied to detect stress-relevant genes in Arabidopsis abiotic stress studies (Sanchez-Munoz et al. 2025) and to refine core gene sets in multi-stress barley meta-analysis (Panahi et al. 2024). The integration of ML with meta-analysis provides complementary discovery framework that enhances sensitivity for genes involved in variable or condition/context-specific stress programs.

Identifying stress-responsive gene sets in common bean is biologically informative, but its practical relevance depends on whether the candidates could translate to improved trait prediction via genomic prediction and if this can be replicated in a closely related species like cowpea. Genomic prediction uses genome-wide marker data to estimate breeding values for any trait, with standard genomic best linear unbiased prediction (GBLUP) model treating all single nucleotide polymorphism (SNP) markers as equally informative (VanRaden 2008). Conversely, assigning higher weights to or considering SNPs in genomic regions with known functional relevance (for example, SNPs proximal to stress-responsive genes) can improve prediction accuracy over equal-weight GBLUP by concentrating genomic signal in regions more likely to harbor causal variants. Common bean and its close relative cowpea (*Vigna unguiculata* L. Walp) share substantial genomic synteny (Lonardi et al., 2019) and orthology-based transfer of functional gene information between species has been applied in legume genomics to leverage gene expression data from one species for marker prioritization in another (Chai et al., 2017). However, to our knowledge, no study to date has connected RNA-seq meta-analysis-derived gene sets to genomic prediction in either common bean or cowpea.

Here, we identified conserved stress-responsive genes in common bean through RNA-seq meta-analysis using Fisher’s combined probability and random-effects restricted maximum likelihood (REML) frameworks and extended the gene set using a support vector machine classifier. Next, we created sets of gene candidates that are functionally linked to stress tolerance and evaluated whether SNP subsets based on gene candidates derived from the stress-tissue meta-analysis and machine learning frameworks improves genomic prediction for stress tolerance in common bean and if this could be extended to cowpea. Our results showed that integrating multiple approaches for meta-analysis and adoption of a suitable machine learning classifier identifies functionally informative stress-response genes. Further, leveraging the candidates identified herein largely improved trait prediction accuracy and uncovered regions that could be targeted for stress resilience breeding in common bean, cowpea and other phylogenetically related leguminous crops.

## Results

### Construction of DEG libraries and cross-study analysis

A total of 19 RNA-seq studies met quality control criteria from 22 initially retrieved datasets, comprising 461 samples and spanning abiotic stress conditions (cold, drought, heat, salt, heavy metal, nitrogen and phosphorus deficiency) and biotic stress conditions like anthracnose, fusarium wilt, halo blight (caused by pathogens including *Fusarium oxysporum*, *Pseudomonas syringae pv phaseolicola*, *Xanthomonas citri pv. fuscans*, *Colletotrichum lindemuthianum*), across four stress-tissue groups: abiotic leaf (n = 7), biotic leaf (n = 6), abiotic root (n = 3), and biotic root (n = 3) (Table S1). Sample counts per study ranged from 4 to 90, and total DEGs per study ranged from 16 to 9,702; per-study DEG counts and contrast summaries are displayed in Fig. 1A. Between 21,731 and 23,856 genes were analyzed per group after expression-based filtering. Hierarchical clustering of cross-study DEG overlap profiles revealed that studies grouped by transcriptional scale rather than stress category in that a high-overlap cluster spanning both abiotic and biotic conditions showed substantially greater pairwise gene overlap than a distinct low-overlap cluster of studies with smaller total DEG sets (Fig. 1B-C).

**Figure 1.**
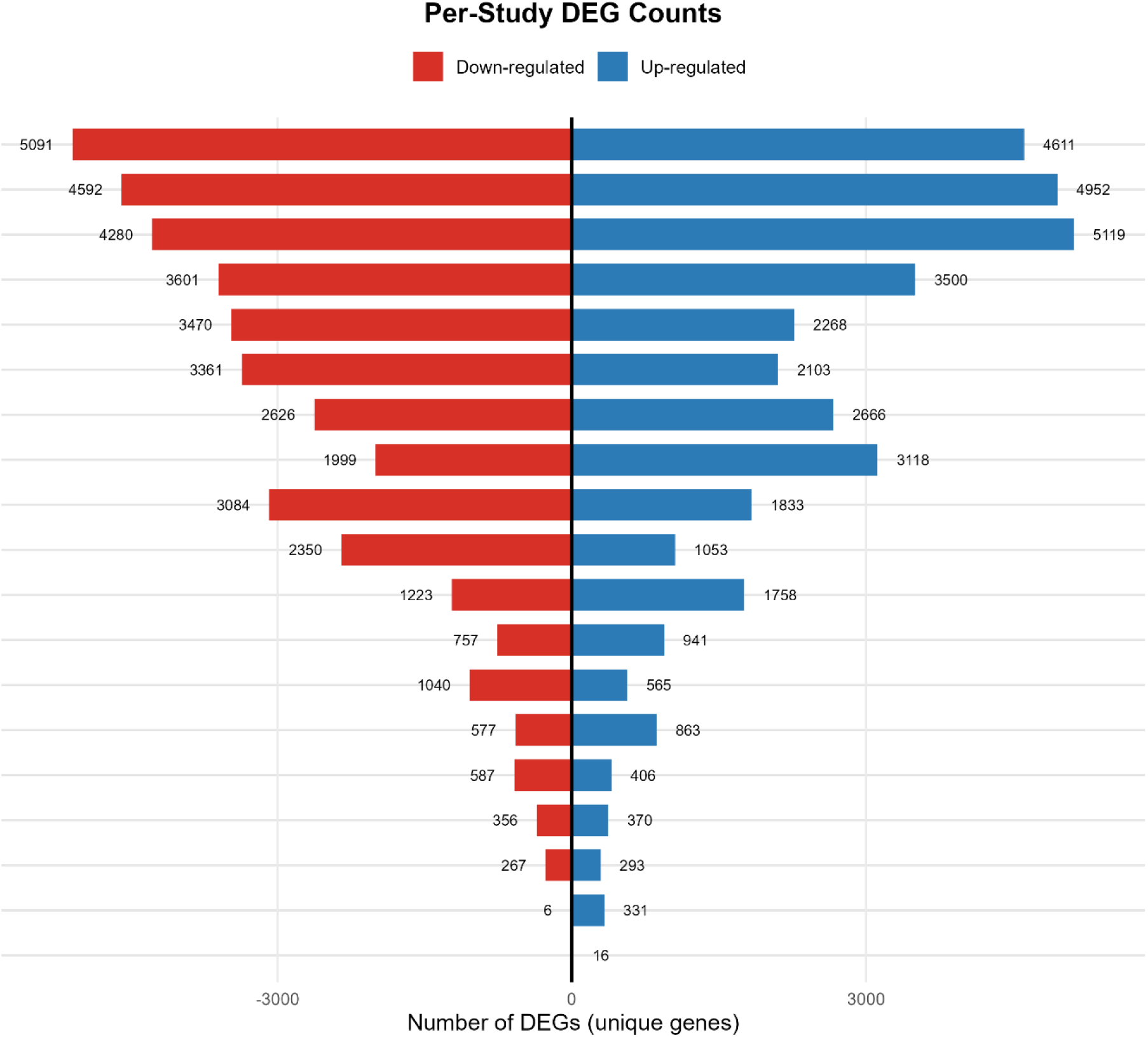
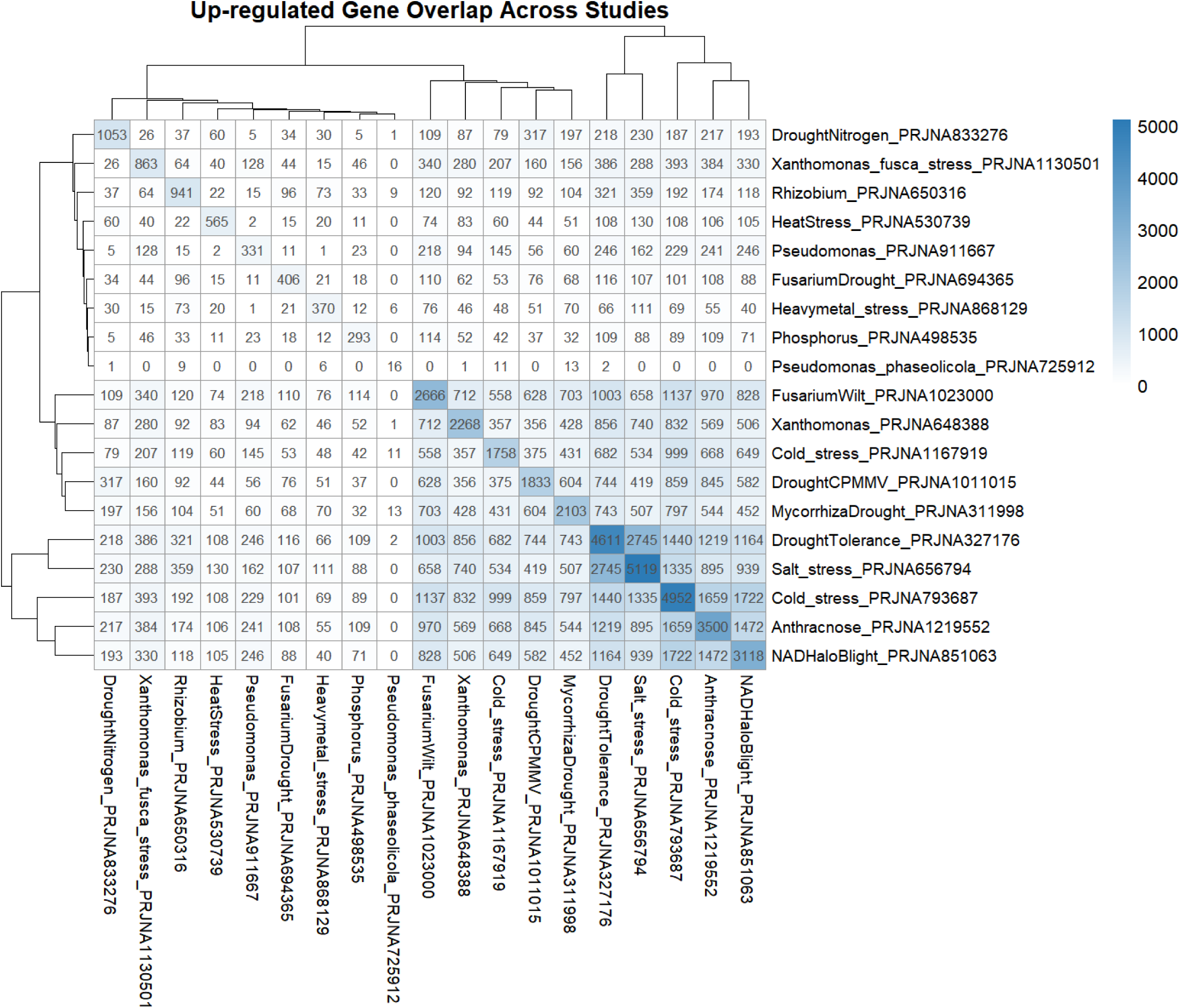
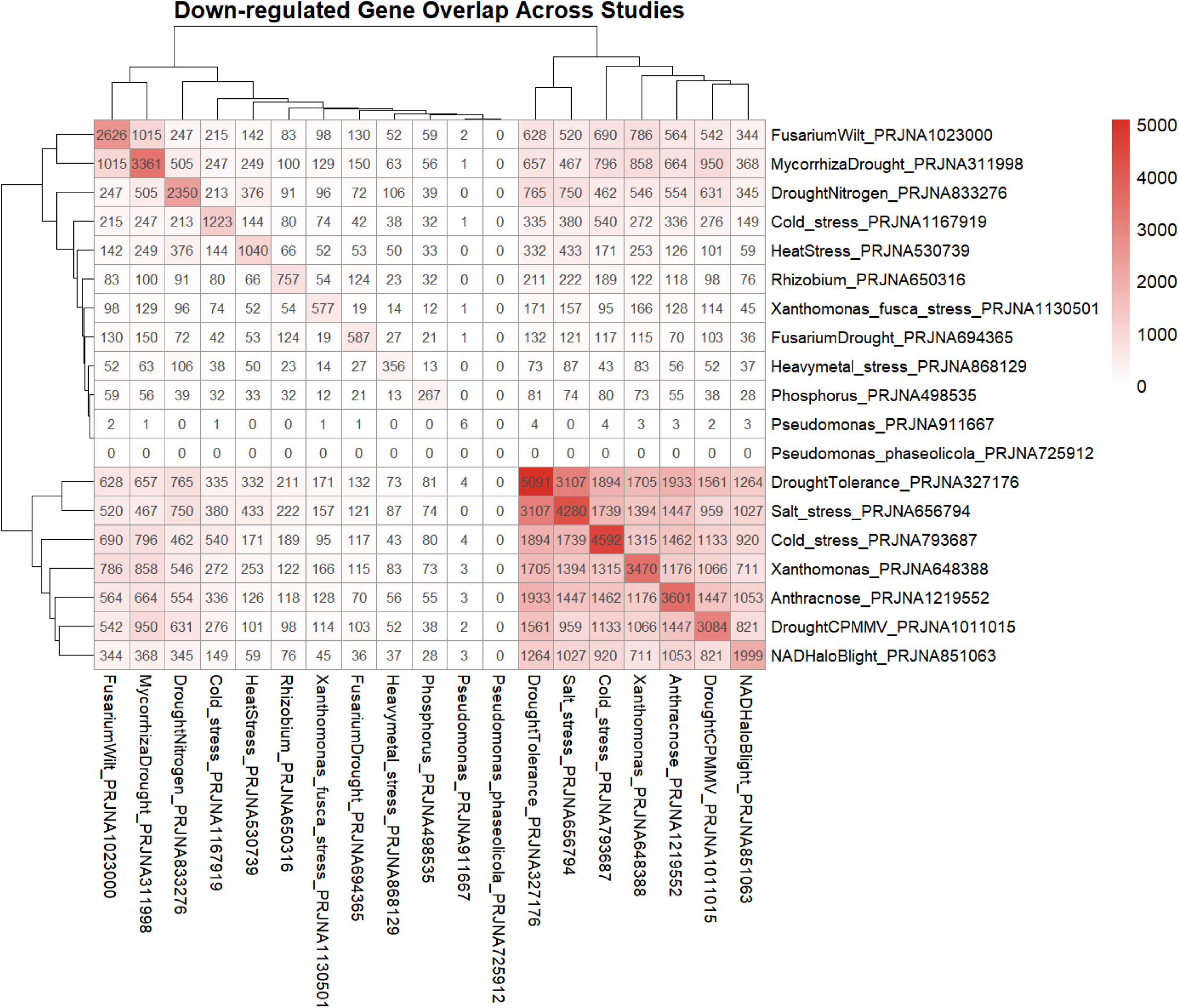
Study-level RNA-seq data quality and cross-study gene expression overlap. (A) Diverging barplot of up- and downregulated DEG counts per study. Jaccard similarity heatmaps showing pairwise overlap of upregulated (B) and downregulated (C) DEGs between the 19 included studies.

### Meta-analysis identifies consensus stress-responsive genes

Consensus meta-DEGs were identified in all four stress-tissue groups using a dual-criterion approach combining Fisher’s combined probability method and REML meta-analysis, with expression direction consistency required for final classification (Table 1; Fig. 2). The abiotic root group yielded the largest consensus meta-DEG set (2,972 genes of 21,731 analyzed; 13.7%), followed by biotic leaf (1,072 of 22,521; 4.8%), abiotic leaf (250 of 23,856; 1.0%), and biotic root (19 of 23,215; 0.1%). In the abiotic leaf group, 222 of 250 consensus meta-DEGs (88.8%) were upregulated, reflecting a predominantly activating transcriptional response to abiotic stress, with 28 genes downregulated. In the abiotic root group, 1,882 genes were upregulated and 1,090 downregulated, indicating broader bidirectional transcriptional reprogramming. The biotic leaf group showed a more balanced directional distribution, with 593 upregulated and 479 downregulated consensus meta-DEGs, and the biotic root group similarly yielded 11 upregulated and 8 downregulated genes across its consensus meta-DEGs. Cross-group overlap of consensus meta-DEGs was minimal, with the largest pairwise intersection between abiotic root and biotic leaf (38 upregulated and 13 downregulated shared genes) and no gene shared across all four groups simultaneously. Permutation tests confirmed that the observed consensus meta-DEG counts exceeded null expectations in all four groups (permutation p < 0.001 for all groups).

**Figure 2.**
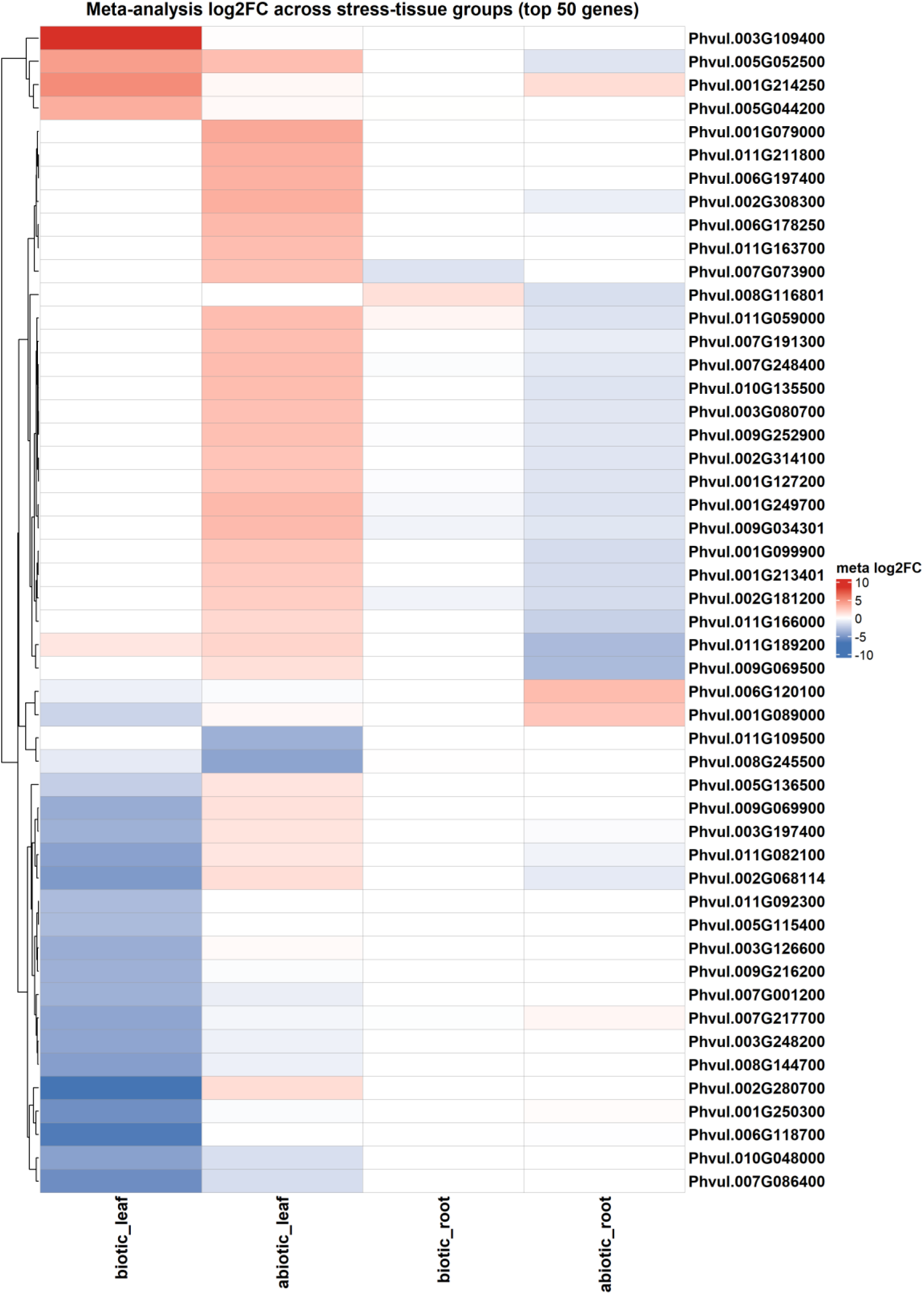
Meta-analysis results across four stress-tissue groups. Heatmap of meta-log_2_FC values for the top 50 most variable consensus meta-DEGs across stress-tissue groups. Red = upregulated; blue = downregulated.

**Table 1.** Summary of differentially expressed genes identified by consensus meta-analysis (meta-DEGs) per stress-tissue group.

| Stress-tissue group | Studies (n) | Genes analyzed | Consensus meta-DEGs | Upregulated | Downregulated | p-value |
| --- | --- | --- | --- | --- | --- | --- |
| Abiotic Leaf | 7 | 23856 | 250 | 222 | 28 | < .001 |
| Abiotic Root | 3 | 21731 | 2972 | 1882 | 1090 | < .001 |
| Biotic Leaf | 6 | 22521 | 1072 | 593 | 479 | < .001 |
| Biotic Root | 3 | 23215 | 19 | 11 | 8 | < .001 |

Genes analyzed = unique genes with quantified expression across all studies in the group. Consensus meta-DEGs = genes significant by both Fisher’s combined probability method and REML meta-analysis at FDR < 0.05 and consistent direction of expression. Upregulated and Downregulated counts refer to the direction of the meta-log_2_FCestimate. p-value = empirical p-value from 1,000 permutations of study labels, testing whether the observed consensus DEG count exceeds the null expectation.

### Machine learning identifies stress-responsive genes beyond consensus meta-DEGs

SVM classification was performed independently for each stress-tissue group (Table 2; Fig. 3). The SVM confirmed a subset of pre-classified genes as ML-confirmed (1 to 1 transitions) and identified additional genes not in the meta-analysis consensus as ML-discovered (0 to 1 transitions). Classification precision ranged from 0.198 (biotic root) to 0.343 (abiotic root), exceeding the null base rate of 0.1699 in all groups (Fig. 3A). Permutation tests confirmed that observed precision exceeded the null distribution in all groups (p < 0.001 for abiotic leaf, abiotic root, and biotic leaf; *p* = 0.02 for biotic root). No gene was universally identified across all four stress-tissue groups (Fig. 3B), and the limited cross-group overlap indicated predominantly stress- and tissue-specific transcriptional programs, with the abiotic root group contributing the largest unique gene set (Table 2).

**Figure 3.**
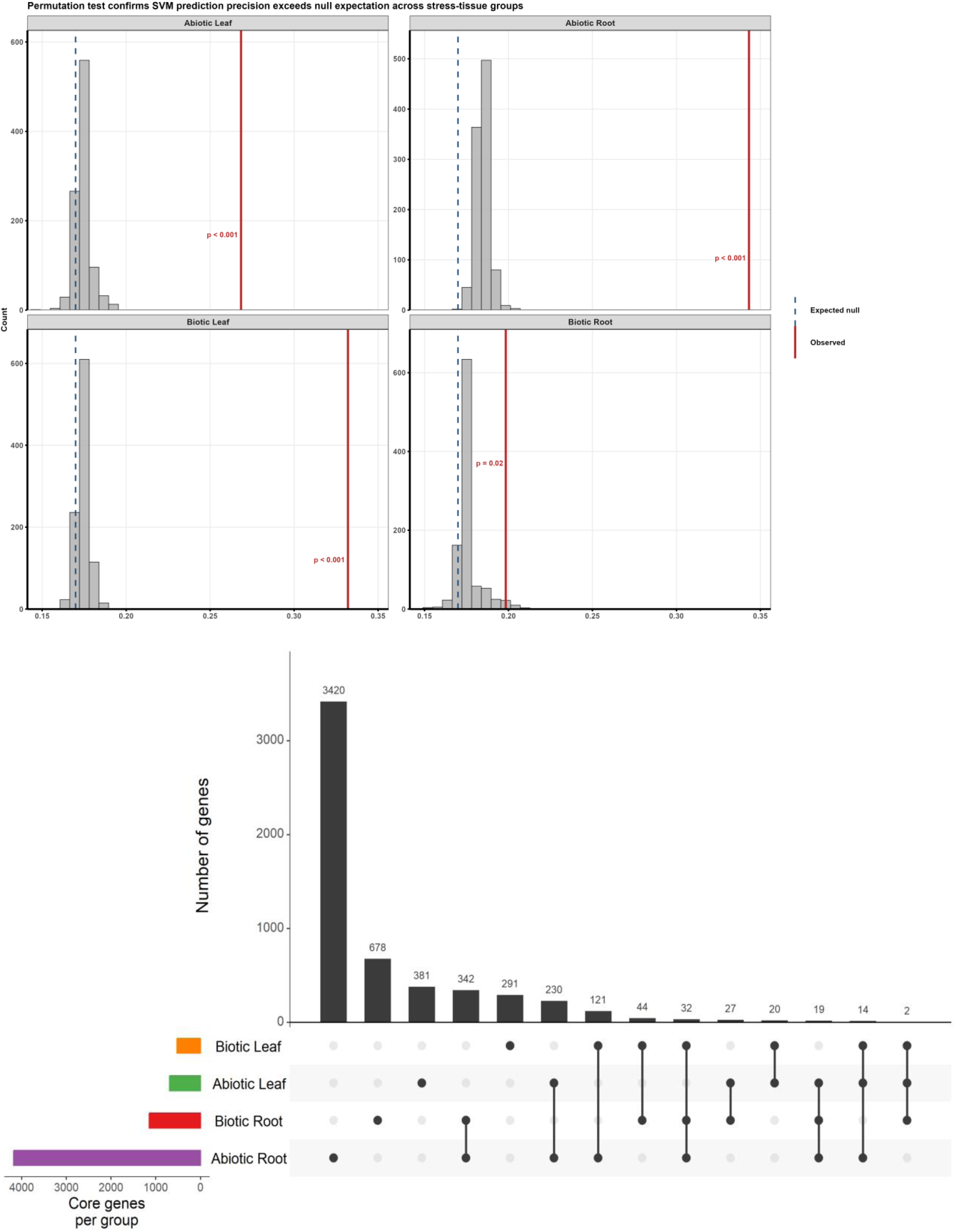
Machine learning (SVM) gene discovery performance. (A) Permutation test confirming that SVM prediction precision exceeds the null expectation (dashed line = null base rate = 0.1699) in all four stress-tissue groups. Distributions represent 1,000 permutations of pre-classification labels; red vertical line = observed precision. (B) UpSet plot of predictor genes (SVM-confirmed and ML-discovered) across stress-tissue groups. No gene was universally identified across all four groups.

**Table 2.** Support vector machine (SVM) classification performance per stress-tissue group.

| Stress-tissue group | Total genes | Pre-classified | Confirmed | ML-discovered | Precision | p-value |
| --- | --- | --- | --- | --- | --- | --- |
| Abiotic Leaf | 23846 | 4134 | 186 | 507 | 0.268 | < .001 |
| Abiotic Root | 21721 | 4012 | 1434 | 2744 | 0.343 | < .001 |
| Biotic Leaf | 22510 | 3917 | 174 | 350 | 0.332 | < .001 |
| Biotic Root | 23186 | 4049 | 227 | 917 | 0.198 | .02 |

Total genes = unique genes used as SVM features; genes with insufficient variance were excluded during SVM feature preparation. Pre-classified = genes labeled as relevant prior to SVM training based on consensus meta-DEG status in any of the four stress-tissue groups. Confirmed = pre-classified genes retained as relevant by SVM (1 to 1 transitions). ML-discovered = genes not pre-classified but identified as relevant by SVM (0 to 1 transitions). Precision = TP/(TP + FP) = Confirmed/(Confirmed + ML-discovered); measures the signal-to-noise ratio of SVM discovery. p-value = empirical p-value from 1,000 label-shuffle permutations testing whether observed precision exceeds the null distribution.

### Weighted gene co-expression network analysis (WGCNA)

WGCNA co-expression networks were constructed for each stress-tissue group using the union of consensus meta-DEGs and ML-identified core genes as input. Co-expression overlap between ML hub genes and consensus meta-DEGs varied markedly across groups: biotic root showed low overlap, indicating that ML hub genes in that group were identified largely independently of the consensus meta-DEG set, whereas biotic leaf showed complete convergence, with hub genes from both sources represented in all co-expression modules (Fig. 4). In the abiotic root group, the turquoise module showed near-equal ML and consensus meta-DEGs representation (27 ML hub genes and 23 consensus hub genes), while the blue module was consensus-exclusive. In the abiotic leaf group, convergence was partial, with the turquoise module ML-dominant, the blue module consensus-dominant, and the brown module consensus-exclusive.

**Figure 4.**
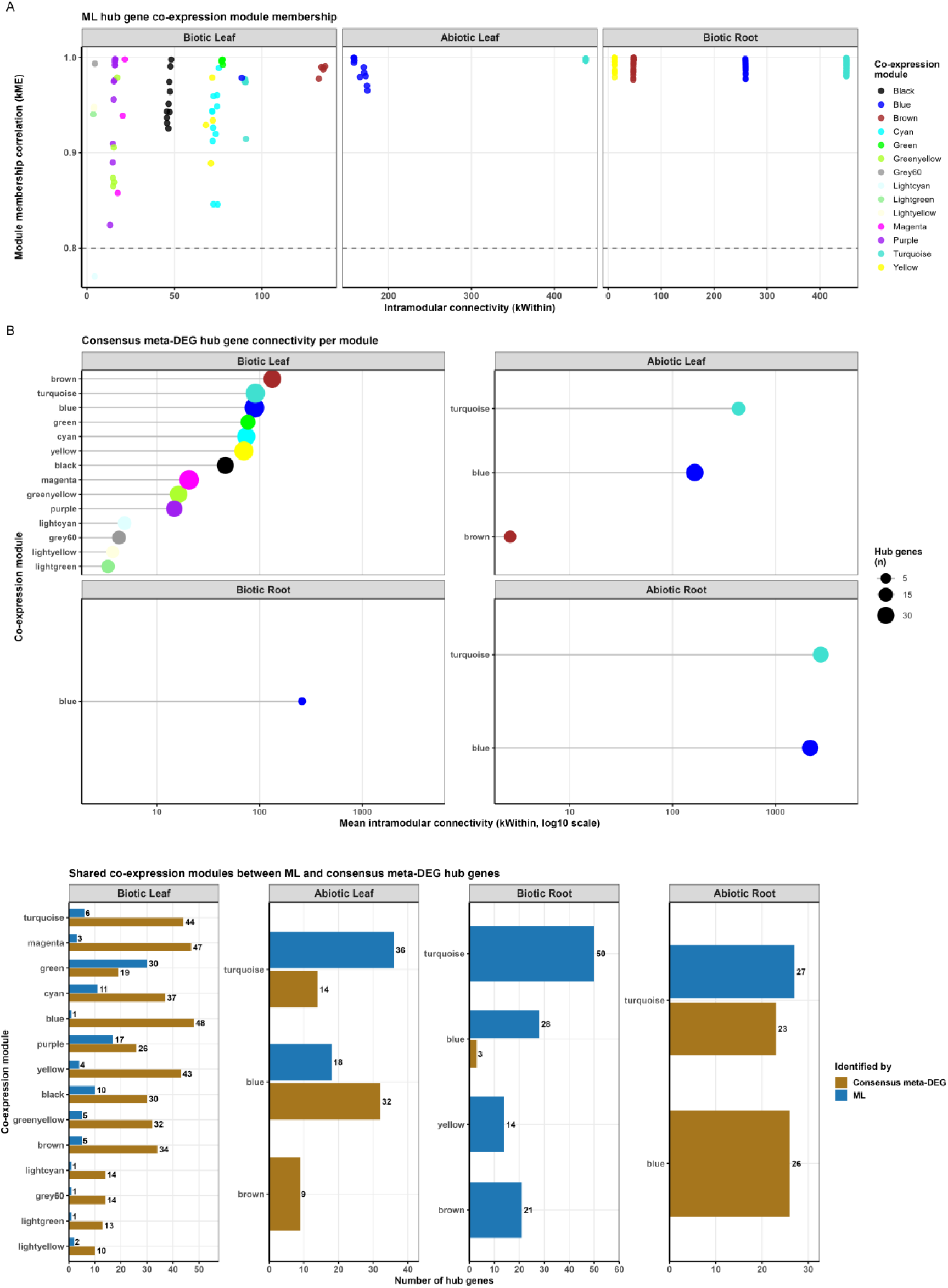
WGCNA co-expression module analysis of hub genes across stress-tissue groups. (A) Module membership (kME) vs. intramodular connectivity (kWithin) for ML-identified hub genes. All hub genes show kME > 0.80 (dashed reference line), confirming strong module membership. Abiotic root contained a single co-expression module (turquoise; kWithin ≈ 2,766, kME = 1.0) with no variation to display and is excluded. (B) Mean intramodular connectivity (kWithin, log10 scale) per consensus meta-DEG co-expression module per group; dot size = number of hub genes per module. (C) Number of hub genes per module per group for ML-identified (blue) and consensus meta-DEG (amber) hub gene sets; bars present in both indicate convergent module identification across discovery approaches.

### Protein-Protein Interaction network of hub genes and gene family identification

PPI subnetworks were constructed from WGCNA hub genes per stress-tissue group and differed substantially in density and biological composition across groups (Fig. 4). The biotic leaf network was the most connected (383 nodes, 1,724 edges), with ribosomal proteins occupying the highest-degree hub positions: four of the top five hubs were ribosomal subunit proteins (large subunit uL6c (Phvul.004G116200), bL19 (Phvul.002G001200), small subunit uS10c [Phvul.001G030600], uS9c [Phvul.001G017500]), indicating that translational machinery components form the primary interaction scaffold in the biotic leaf stress response; a mitogen-activated protein kinase, (MAPK; Phvul.003G059500) and a MAP kinase kinase (Phvul.010G163000) also ranked among the top hubs, consistent with MAPK cascade activation as a central signal transduction mechanism in defense signaling, and all 12 biotic leaf hubs were consensus meta-DEGs. In the abiotic leaf network (59 nodes, 46 edges), transcription factors were the dominant hub class (7 of 20), including AP2/ERF domain (Phvul.004G068900), growth-regulating factor (Phvul.010G130000), and bHLH domain-containing proteins (Phvul.001G026800), reflecting a transcription factor-centered regulatory scaffold under abiotic leaf stress; a tubulin beta chain (Phvul.002G198600) was the highest-connectivity node, identifying cytoskeletal remodeling as an additional network focal point. The abiotic root network (52 nodes, 32 edges) was anchored by hormone signaling components: NPR1-like protein (Phvul.006G131400), a salicylic acid signaling regulator, ranked joint-highest in connectivity (tied with a fungal lipase-like domain-containing protein, Phvul.004G020000), and COI1 (Phvul.008G226500), the F-box component of the SCF^COI1^ jasmonate receptor complex, was among the prominent signaling hubs, indicating that SA-JA hormone pathway genes occupy central positions in the abiotic root interaction landscape; heat shock protein 90 (Phvul.004G107700), an ML-discovered gene, was also among the top hubs. The biotic root network was extremely sparse (26 nodes, 6 edges), with all 12 hub genes derived from ML-discovered genes and hormone signaling genes comprising the largest functional class (6 of 12), including a chitinase (Phvul.001G046400), a RING-type domain-containing protein (Phvul.002G099600), and a prohibitin (Phvul.011G018300). No gene was identified as a PPI hub in more than one stress-tissue group, and hub gene meta-log_2_FCprofiles across all four groups are displayed in Fig. 5; 6.

**Figure 5.**
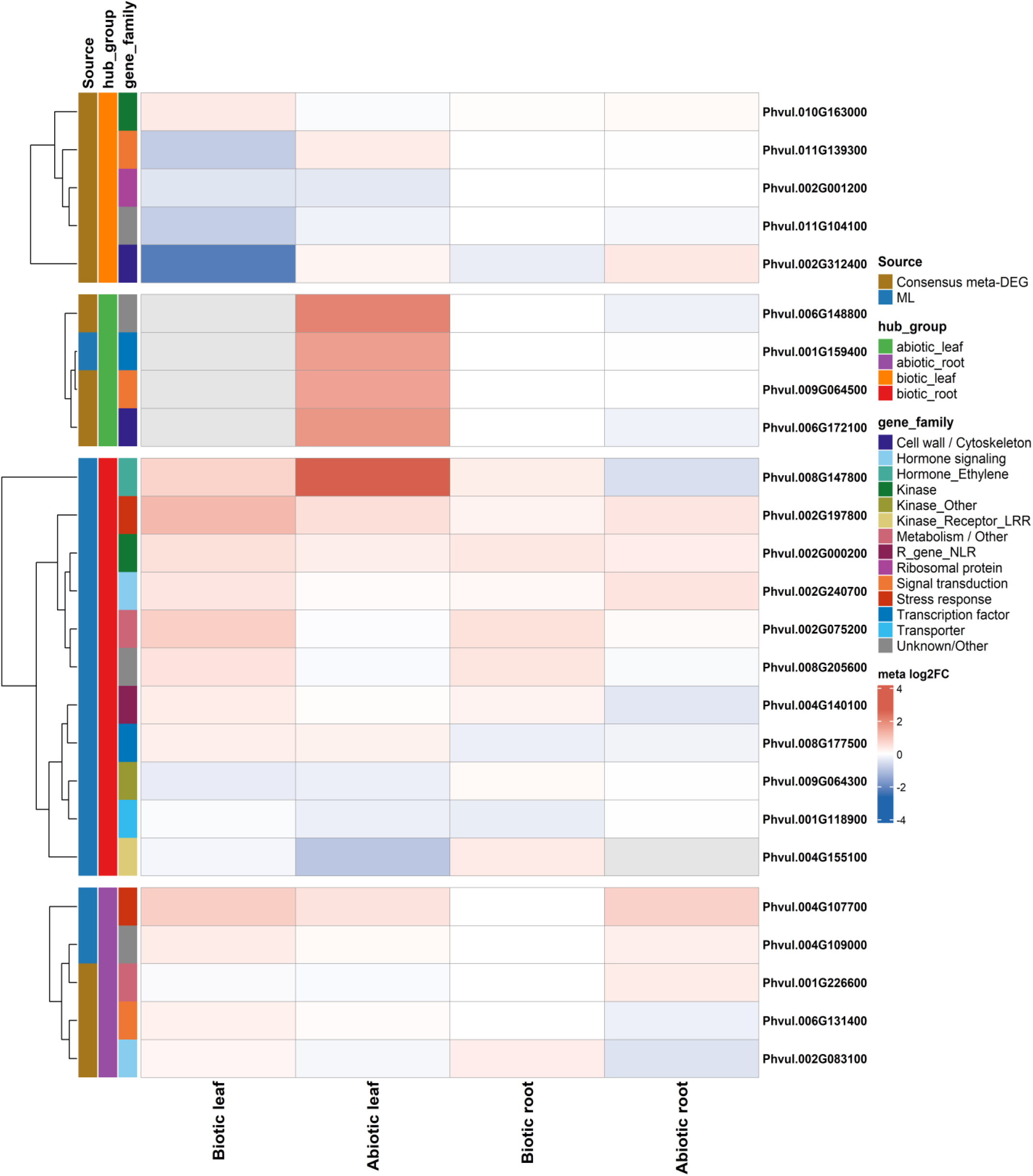
Hub gene expression heatmap across stress-tissue groups. Meta-log2FC expression values for the top 12 PPI hub genes per stress-tissue group (48 genes total). Rows split by the PPI network in which each gene was identified as a hub. Annotation bars indicate gene source (consensus meta-DEG = amber; ML-discovered = blue; both = charcoal) and functional gene family.

**Figure 6.**
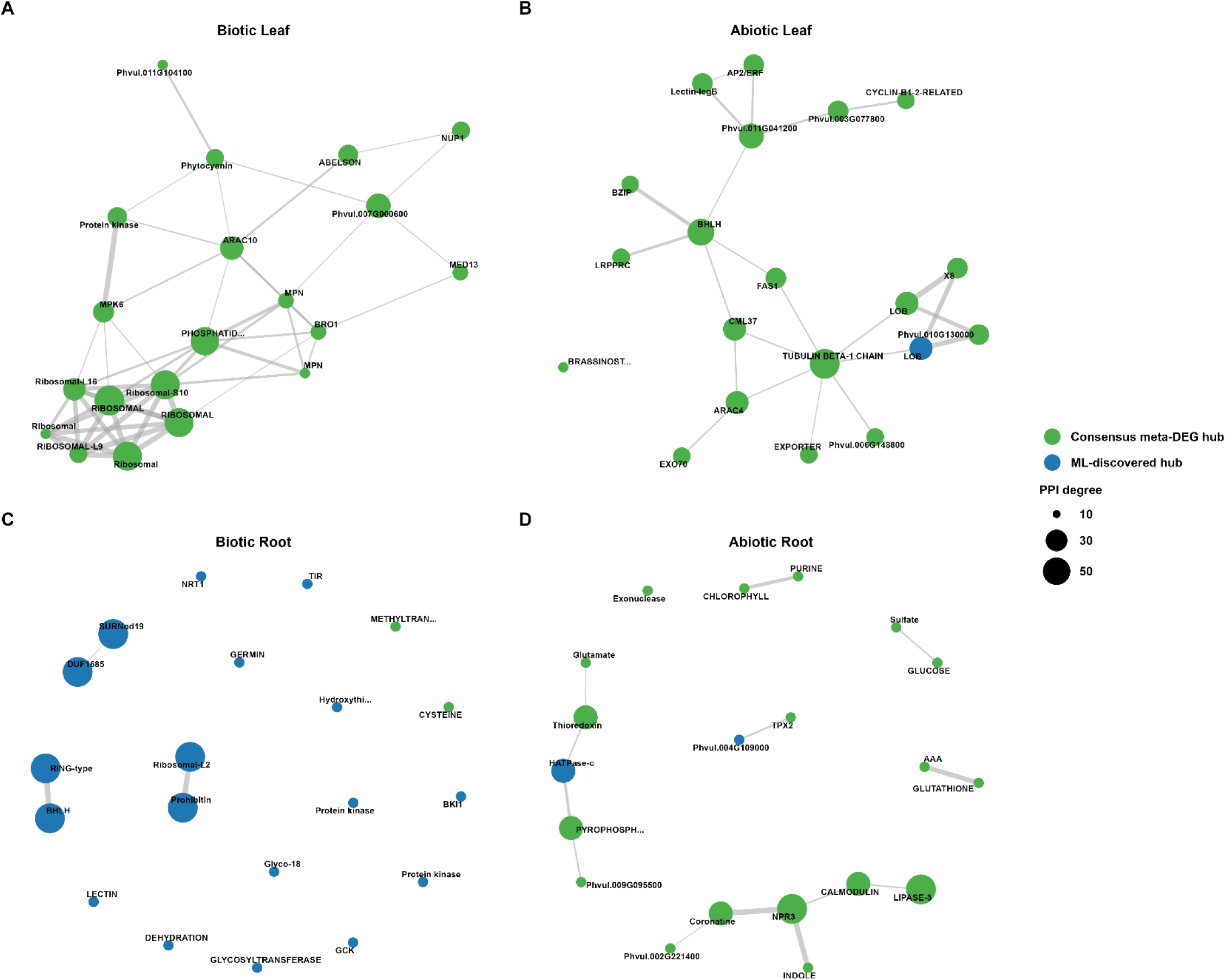
Protein-protein interaction (PPI) networks of hub genes across stress-tissue groups. Networks constructed using STRINGdb (minimum interaction score = 200). Top 20 hub nodes per panel shown; edges filtered to hub-to-hub interactions. Node color: green = consensus meta-DEG hub; blue = ML-discovered hub. Node size proportional to PPI degree. Edge width proportional to interaction confidence score.

### Functional annotation of hub and other genes

Hub genes were annotated against GO, KEGG, Pfam, and Panther databases and classified into seven functional gene family categories (Table 3). Hormone signaling genes constituted the largest category (105 hub gene-group combinations), followed by Other/Unknown (98), kinases (27), transcription factors (26), stress response proteins (19), transporters and cell wall genes (10), and disease resistance (R) genes (6). The biotic root group had the highest proportion of hormone signaling genes (45 of 113; 39.8%), consistent with jasmonate and ethylene signaling roles in root defense. Disease resistance R genes were identified in the biotic leaf, abiotic leaf, and biotic root groups (2 per group), with none detected in abiotic root.

**Table 3.** Distribution of WGCNA hub genes across functional gene family categories and stress-tissue groups.

| Gene family | Biotic leaf | Abiotic leaf | Biotic root | Abiotic root | Total |
| --- | --- | --- | --- | --- | --- |
| Hormone signaling | 34 | 17 | 45 | 9 | 105 |
| Kinases | 5 | 5 | 13 | 4 | 27 |
| Transcription factors | 8 | 6 | 7 | 5 | 26 |
| Stress response proteins | 9 | 5 | 4 | 1 | 19 |
| Transporters and cell wall | 2 | 2 | 5 | 1 | 10 |
| Disease resistance (R genes) | 2 | 2 | 2 | 0 | 6 |
| Other/Unknown | 37 | 17 | 37 | 7 | 98 |

Hub genes defined as the top 50 most-connected nodes by intramodular connectivity (kWithin) per WGCNA co-expression module, restricted to ML-discovered genes; all hub genes showed kME > 0.80, confirming strong module membership. Thirty detailed sub-categories condensed into seven broad functional families. Other/Unknown = genes with no assignable Pfam, Panther, or KEGG Orthology (KO) annotation. Total = sum across all four stress-tissue groups; a gene appearing as a hub in multiple groups is counted once per group.

Functional enrichment analysis of consensus meta-DEGs per stress-tissue group identified significant GO and KEGG enrichment exclusively in the abiotic leaf group; no other group yielded statistically significant results after FDR correction (Table 4). Enriched GO terms included peroxidase activity (GO:0004601; 7 genes; fold enrichment [FE] = 4.00; FDR = 0.031), response to oxidative stress (GO:0006979; 7 genes; FE = 3.96; FDR = 0.031), root development (GO:0048364; 3 genes; FE = 7.90; FDR = 0.038), shoot system development (GO:0048367; 3 genes; FE = 7.90; FDR = 0.038), and enzyme inhibitor activity (GO:0004857; 6 genes; FE = 3.67; FDR = 0.038). Enriched KEGG orthologous groups included carbon-nitrogen hydrolase (K01674; 2 genes; FE = 9.63; FDR = 0.045), WRKY transcription factor (K04125; 3 genes; FE = 7.78; FDR = 0.045), and peroxidase (K00430; 7 genes; FE = 2.71; FDR = 0.045). The convergence of peroxidase enrichment across both GO (GO:0004601, GO:0006979) and KEGG (K00430) analyses provides independent confirmation of oxidative stress defense as a central feature of the abiotic leaf stress response.

**Table 4.** Functional enrichment of WGCNA hub genes in the abiotic leaf stress-tissue group.

| Type | ID | Term | Ontology | Gene count | Fold enrichment | FDR |
| --- | --- | --- | --- | --- | --- | --- |
| GO | GO:0006979 | response to oxidative stress | Biological Process | 7 | 3.96 | 0.0307 |
| GO | GO:0004601 | peroxidase activity | Molecular Function | 7 | 4.00 | 0.0307 |
| GO | GO:0048364 | root development | Biological Process | 3 | 7.90 | 0.0383 |
| GO | GO:0048367 | shoot system development | Biological Process | 3 | 7.90 | 0.0383 |
| GO | GO:0004857 | enzyme inhibitor activity | Molecular Function | 6 | 3.67 | 0.0383 |
| KEGG | K01674 | Carbon-nitrogen hydrolase | KEGG Orthology | 2 | 9.63 | 0.0445 |
| KEGG | K04125 | WRKY transcription factor | KEGG Orthology | 3 | 7.78 | 0.0445 |
| KEGG | K00430 | Peroxidase | KEGG Orthology | 7 | 2.71 | 0.0445 |

Only the abiotic leaf group yielded significant enrichment after false discovery rate (FDR) correction (Benjamini-Hochberg). GO enrichment tested using Fisher’s exact test against the *P. vulgaris* genome annotation background. KEGG enrichment tested against KO assignments. Fold enrichment = observed/expected gene count ratio. No significant terms were identified for biotic leaf, abiotic root, or biotic root groups after FDR correction (minimum FDR = 0.050 for abiotic root; excluded). KO descriptions for K00430, K01674, and K04125 are based on KEGG Orthology definitions.

### Independent drought experiment

Physiological measurements confirmed differential drought tolerance between the two contrasting common bean genotypes across two independent greenhouse trials. SPAD-derived chlorophyll content declined under water-deficit conditions in both genotypes, with a significant genotype × day interaction (split-plot repeated-measures ANOVA, α = 0.05) and a decline in the susceptible genotype (PI361408) becoming apparent from day 6 of water stress in Trial 1. Mean chlorophyll values under drought were consistently higher in the tolerant genotype (PI549793) than in the susceptible genotype in both trials (Trial 1: 28.1 vs. 23.0; Trial 2: 37.3 vs. 33.6), whereas values under well-watered conditions were comparable between genotypes (Trial 1: 31.8 vs. 29.4; Trial 2: 39.7 vs. 39.6), indicating a stress-specific rather than constitutive difference in chlorophyll retention (Table S3). Leaf relative water content did not differ significantly between genotypes overall, although the susceptible genotype showed a modest increase in relative water content at 8 and 10 days after stress onset, likely reflecting reduced transpirational water loss associated with advanced leaf senescence rather than improved drought tolerance (Fig. S4 - S8). Reproductive and survival phenotypes showed the clearest genotypic divergence: flowering (28 days after planting) and pod emergence (34 days after planting) occurred exclusively in the tolerant genotype, with no flowering or pod set observed in the susceptible genotype. At 20 days after water withdrawal, plant mortality reached 70% in the tolerant genotype compared with 100% in the susceptible genotype.

Differential expression analysis of the independent common bean drought experiment identified 1,848 DEGs at T3 (144 hours post-stress initiation) and 1,318 DEGs at T4 (192 hours). The intersection of the T3 and T4 DEG sets defined 509 sustained drought-responsive genes. and permutation testing of condition labels (1,000 permutations) confirmed that this count substantially exceeded the null expectation, as all 1,000 permutations yielded fewer sustained genes than the observed count (maximum permuted count = 281; empirical p < 0.001). Genotype-stratified analysis identified 1888 and 2722 significant genes in the tolerant (PI549793) and susceptible (PI361408) genotypes, respectively, at T3, and 1,459 and 1,662 genes at T4, with the susceptible genotype showing a consistently larger and more statistically extreme transcriptional response at both timepoints (Figure S9).

KEGG pathway enrichment of the 509 sustained drought-responsive genes identified 5 enriched pathways, with plant hormone signal transduction (ko04075; 7 of 30 KO-annotated sustained genes; fold enrichment [FE] = 53.8; FDR < 0.001), MAPK signaling pathway (ko04016; 5 genes; FE = 44.9; FDR < 0.001), plant-pathogen interaction (ko04626; 4 genes; FE = 22.3; FDR < 0.001), phenylpropanoid biosynthesis (ko00940; 3 genes; FE = 44.0; FDR < 0.001), and cutin, suberine and wax biosynthesis (ko00073; 2 genes; FE = 48.5; FDR = 0.005). GO enrichment of the same gene set using a custom TERM2GENE table from the Phytozome v2.1 annotation identified 5 significant term groups, including acyltransferase and biosynthetic process activity (8 genes; FE = 27.5; padj < 0.001) and defense response, receptor signaling, and protein kinase activity (9 genes; FE = 21.5; padj < 0.001). Enrichment of the genotype-stratified gene sets identified plant hormone signal transduction as the most significantly enriched KEGG pathway across all three sets: tolerant-specific (12 of 57 KO-annotated genes; FE = 48.6; FDR < 0.001; 6 pathways total), susceptible-specific (20 of 83 KO-annotated genes; FE = 55.6; FDR < 0.001; 8 pathways total), and shared (9 of 61 KO-annotated genes; FE = 34.0; FDR < 0.001; 10 pathways total), with MAPK signaling and phenylpropanoid biosynthesis also enriched in multiple genotype sets, indicating that hormone signaling cascade gene activation was a consistent feature of the drought transcriptional response irrespective of genotypic tolerance status.

### External validation of drought-responsive gene predictions

Core drought-responsive genes were validated against an independent common bean drought experiment not included in the meta-analysis training set (Fig. 7A). The overlap between SVM core genes and external DEGs was significantly greater than expected by chance (Odds Ratio = 1.35; *p* = .001, Fisher’s exact test). Predicted drought-response probability scores were significantly higher for external DEGs than for non-significant genes (Fig. 7B), indicating that Random Forest probabilities trained on the meta-analysis gene set are likely informative for independent experiment classification. Leave-one-study-out cross-validation (LOSO-CV) across the four drought studies further evaluated model transferability (Table S2).

**Figure 7.**
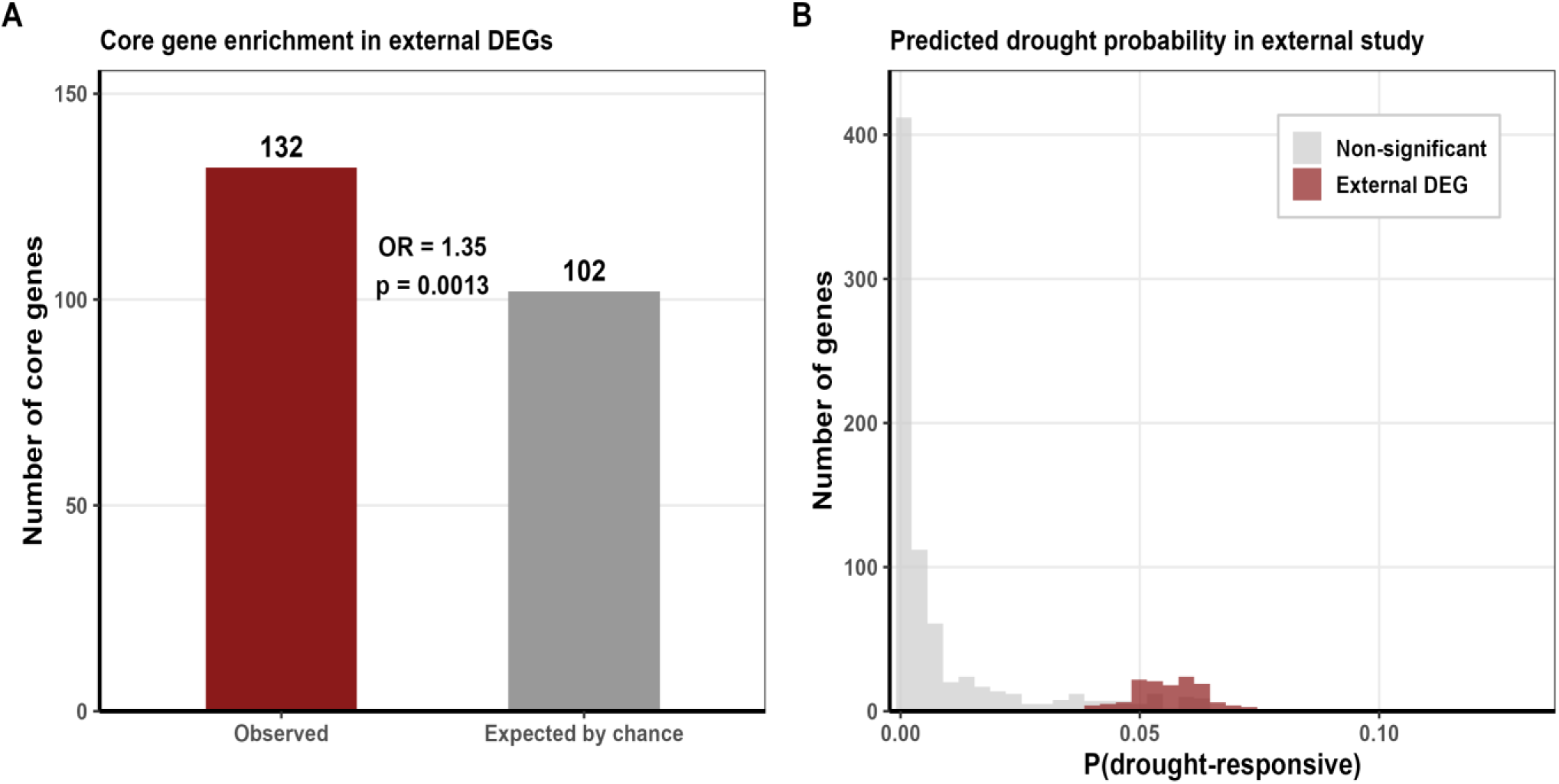
External validation of drought-responsive gene predictions in an independent common bean experiment. (A) Observed (n = 132) vs. expected by chance (n=102) overlap between SVM core drought-responsive genes and DEGs from an independent common bean drought experiment (Odds Ratio = 1.35; *p* = .001, Fisher’s exact test). (B) Distribution of predicted drought-response probabilities for the 902 core genes found in the external dataset, stratified by external DEG classification (firebrick = external DEG; grey = non-significant).

### Ortholog mapping of stress-responsive genes to cowpea

Orthology mapping via Ensembl Plants BioMart identified one-to-one *V. unguiculata* orthologs for 17,779 *P. vulgaris* genes across the abiotic and biotic stress tier classifications. Within the abiotic stress framework, 2,399 cowpea ortholog assignments corresponded to Tier 1 meta-analysis consensus DEGs (mean sequence identity = 90.0%), 2,118 to Tier 3 ML-discovered genes (88.1%), and 53 to Tier 2 ML hub genes (88.0%); the corresponding biotic stress counts were 3,161 (Tier 1; 89.9%), 2,487 (Tier 3; 88.0%), and 185 (Tier 2; 87.0%). Of the 291 ML hub genes (Tier 2) identified across all stress-tissue groups, 238 (81.8%) - comprising 53 from abiotic and 185 from biotic stress groups - had confirmed cowpea orthologs. Functional annotation of these 238 cowpea hub gene orthologs identified Other/Unknown as the predominant class (165 genes; 69.3%), followed by Kinase_Signaling (30; 12.6%), Transcription_Factor (26; 10.9%), Transporter (16; 6.7%), and R_gene_Defense (1; 0.4%); the proportional distribution of functional classes was substantially conserved between *P. vulgaris* hub genes and their cowpea orthologs, with gene-level functional class assignments across both species detailed in the heatmap (Fig. 8).

**Figure 8.**
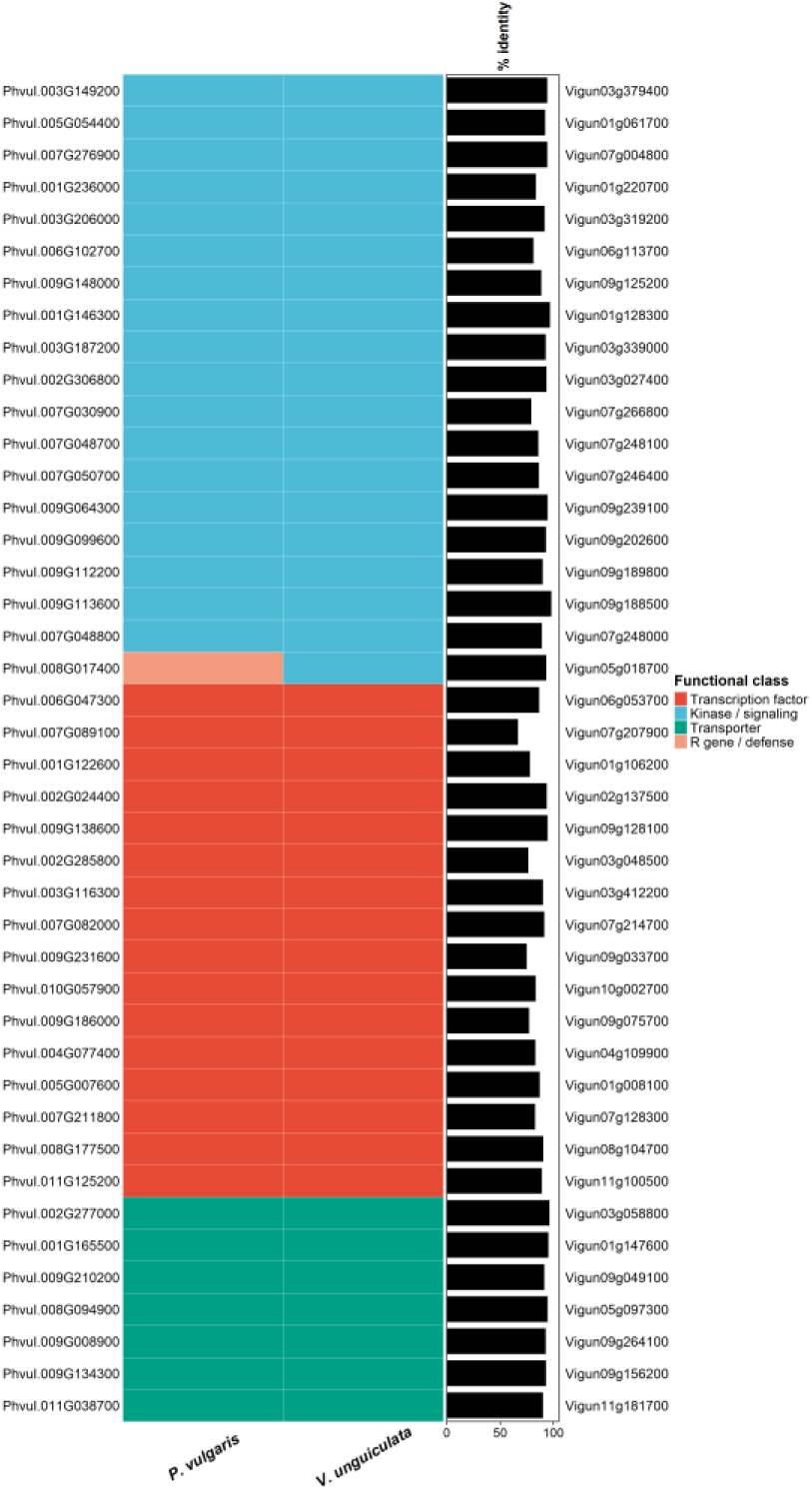
Gene-level functional class assignments of *P. vulgaris* ML hub gene orthologs in cowpea (*Vigna unguiculata*). Heatmap of functional class assignments for non-Other/Unknown *P. vulgaris* ML hub genes with confirmed cowpea orthologs. Each row is one hub gene; left column = *P. vulgaris* gene ID; centre tiles = functional class in *P. vulgaris* (left tile) and *V. unguiculata* (right tile); right bar = percent sequence identity between orthologs (black).

### Genomic prediction using RNA-seq-informed SNP subset scheme

SNP subset schemes based on proximity to evidence gene sets (Meta-DEG, ML genes, Background) were evaluated against standard GBLUP in common bean disease resistance traits and cowpea abiotic stress traits. In common bean (Fig. 9), Meta-DEG SNP subset improved SCN resistance prediction over baseline for all the three HG types (+0.7% to +1.2%), while ML-Combined subset improved prediction for HG Type 7 (+0.3%) and HG Type 2.5.7 (+0.5%) but fell below baseline for HG Type 1.3.6.7 (−0.7%), indicating that the ML hub gene SNP set is less informative than the consensus Meta-DEG set for the most accurately predicted HG type. The Background SNP subset underperformed baseline for all three HG types, confirming that prediction gains were attributable to evidence gene-proximal SNPs rather than to reduced SNP density. The Meta-DEG proximal SNPs exceeded the mean accuracy of 100 size-matched random SNP draws for all three HG types, while the ML-Combined proximal SNPs did not for HG Type 1.3.6.7, consistent with the lower informativeness of the ML-Combined SNP set for that HG type (Fig. 9). For bacterial wilt, Meta-DEG subset outperformed baseline for Race 528 (+0.6%) and Race 557 (+3.3%), while all evidence-based scheme subset modestly underperformed baseline for Race 597 (−1.3%). The Meta-DEG proximal SNPs exceeded the random draw mean for Race 528 and Race 557 but not Race 597, indicating that the prediction advantage of evidence-based SNP selection is race-specific. The consistent ranking of Meta-DEG and ML gene schemes above Background for most disease resistance traits indicated that stress-responsive gene-proximal SNPs carry predictive information relevant to biotic stress resistance, even when derived from gene expression profiling across diverse stress conditions.

**Figure 9.**
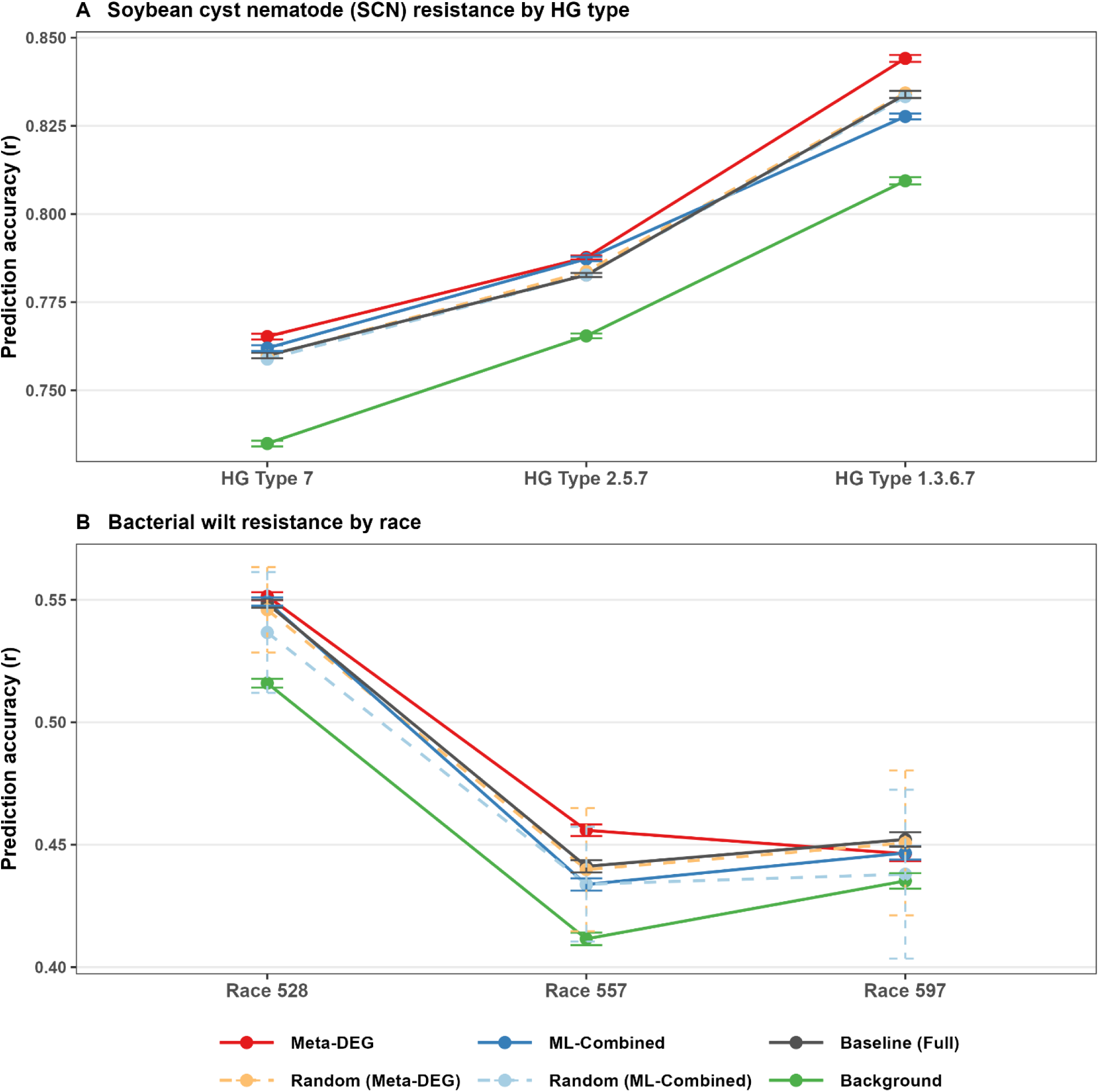
Genomic prediction accuracy for common bean (*Phaseolus vulgaris*) disease resistance traits using RNA-seq-informed SNP subset. (A) Soybean cyst nematode (SCN) resistance across three HG types (B) Bacterial wilt resistance across three races. Meta-DEG = SNPs in/near consensus meta-DEG genes; ML-Combined = SNPs in/near ML-discovered genes; Baseline (Full) = equal-weight full-panel GBLUP; Background = remaining SNPs after evidence gene removal; Random (Meta-DEG) and Random (ML-Combined) = size-matched random SNP draws.

In cowpea (Fig. 10), the ML gene SNP subset achieved the highest prediction accuracy for five of six traits evaluated, with the largest relative gains for the salt stress plant height tolerance index (RTI_H; +86.7%) and fresh biomass tolerance index (RTI_FB; +13.5%); improvements for the remaining drought and salt traits were smaller (0.9%–3.1%). The exception was the composite drought tolerance score (dt_Score), where Meta-DEG SNP subset performed best (+5.4%) and ML gene subset underperformed the baseline (−1.4%). Background SNP schemes performed close to or below baseline across all traits, indicating that prediction gains were attributable to evidence gene-proximal SNPs rather than to differences in SNP density. ML gene-proximal SNPs exceeded the mean accuracy of 100 size-matched random SNP draws in five of six traits (all except dt_Score), while Meta-DEG proximal SNPs exceeded random in only two of six (dt_RTI_C_T and dt_Score), confirming that the refined ML hub gene set carries greater genomic prediction utility for cowpea abiotic stress than the broader consensus meta-DEG set.

**Figure 10.**
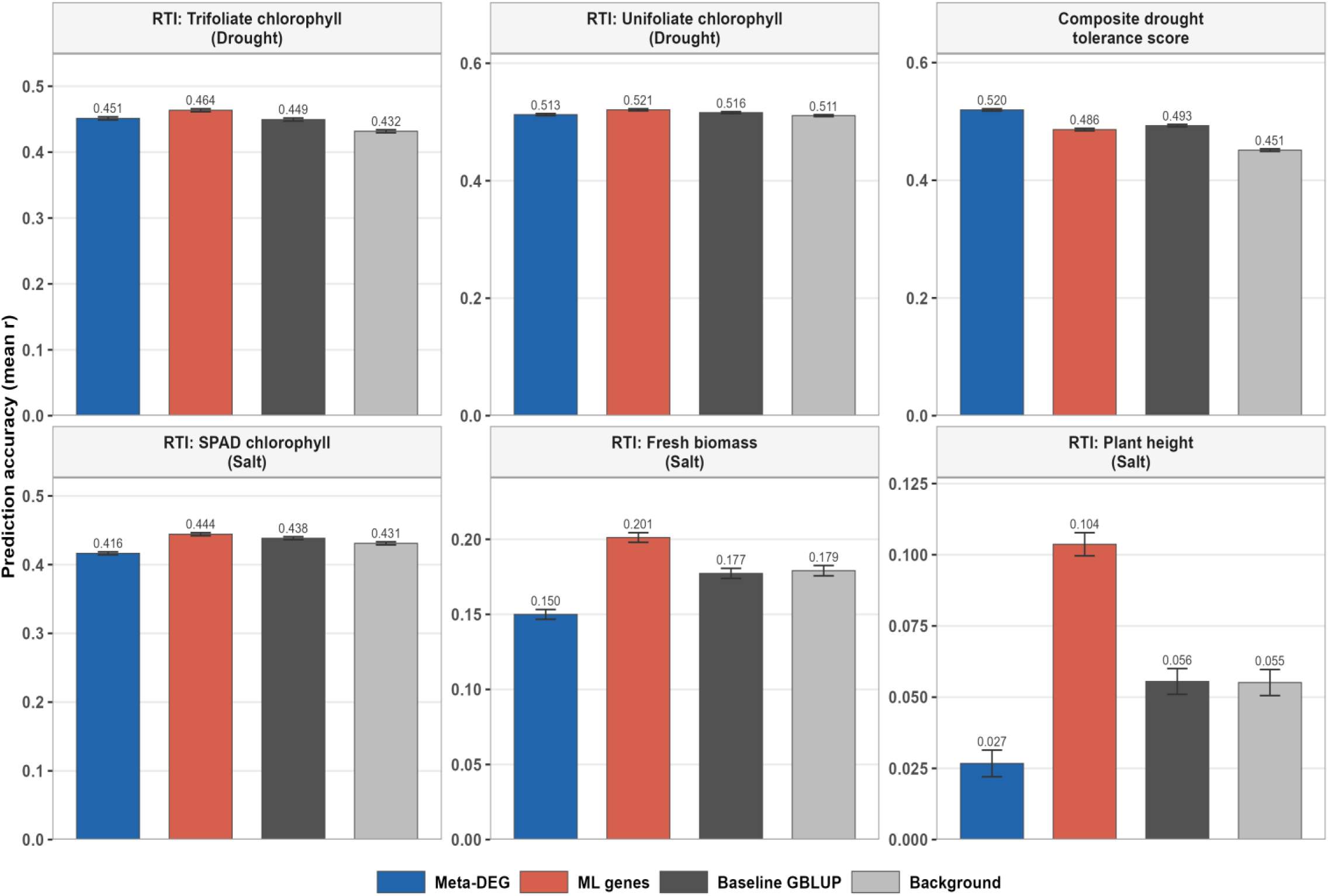
Genomic prediction accuracy for cowpea (*Vigna unguiculata*) abiotic stress traits using RNA-seq-informed SNP subset. Meta-DEG = SNPs in/near consensus meta-DEG genes; ML genes = SNPs in/near ML-discovered genes; Baseline GBLUP = equal SNP weights (standard GBLUP); Background = remaining SNPs after evidence gene removal. RTI = relative tolerance index. Drought traits: RTI for trifoliate chlorophyll (RTI_C_T), unifoliate chlorophyll (RTI_C_U), and composite tolerance score. Salt traits: SPAD chlorophyll (RTI_C), fresh biomass (RTI_FB), and plant height (RTI_H).

## Discussion

### Multi-stress RNA-seq meta-analysis reveals tissue- and condition-specific consensus transcriptomes in common bean

In this study, our dual-criterion meta-analysis framework identified consensus stress-responsive genes independently within each of the four stress-tissue groups, allowing us to directly compare how tissue type and stress category shape the size and composition of the resulting gene sets. Essentially, we found that the size and directionality of consensus meta-DEG sets varied substantially across stress-tissue groups, reflecting genuine biological specificity in tissue responses to stress categories rather than statistical artifact (Pereira et al., 2020). Abiotic root produced the largest consensus set (2,972 genes; 13.7% of analyzed genes), consistent with roots’ central role in abiotic stress perception and water/nutrient uptake (Aroca et al., 2012), which demands broad and coordinated transcriptional reprogramming. In contrast, the abiotic leaf response was comparatively narrow (250 genes; 1.0%), likely because the seven constituent studies spanned diverse abiotic stressors (cold, drought, heat, and salt), producing partially non-overlapping gene sets such that only the most broadly conserved stress-responsive genes passed our dual-criterion consensus filter. The stratified approach we applied, which combines Fisher’s method and REML meta-analysis with heterogeneity-dependent thresholds, is more robust than single-criterion approaches, and we interpreted the narrow abiotic leaf set as reflecting genuine biological constraint rather than insufficient statistical power. Biotic root yielded the fewest consensus meta-DEGs of any group (19 genes), indicating that root defense responses to Fusarium wilt, Rhizobium inoculation, and combined fungal-drought stresses are not transcriptionally uniform across studies; these conditions likely triggered mechanistically distinct programs, defense-oriented for Fusarium versus mutualistic for Rhizobium (Gourion et al., 2015). Permutation testing confirmed that all four consensus counts exceeded the null expectation under 1,000 label permutations (p < 0.001 for all groups), demonstrating that these cross-study consensus signals are not attributable to chance.

### Machine learning extend consensus framework coverage and identifies additional stress-responsive genes

The SVM classification step recovered a substantially larger set of stress-relevant genes in all four stress-tissue groups, and we found that its complementarity to consensus meta-analysis was highest precisely where the consensus set was smallest. The proportion of ML-discovered genes (0 to 1 transitions) was highest in biotic root (917 of 1,144 total core genes; 80.1%) and abiotic root (2,744 of 4,178; 65.6%), showing that SVM-based discovery extracted discriminative patterns from the expression matrix even when between-study heterogeneity prevented deep coverage by consensus approach. By contrast, ML-confirmed genes (1 to 1 transitions) made up a higher proportion of the abiotic leaf core gene set, where the consensus meta-DEG set was already biologically refined. This inverse relationship between consensus set size and ML-discovery proportion indicates that the two methods are functionally complementary rather than redundant. Essentially, meta-analysis captured genes with consistent cross-study evidence, while SVM captured genes with discriminative expression patterns not resolved by the consensus filter. Sanchez-Munoz et al. (2025) used SVM clustering in Arabidopsis abiotic stress data to identify gene modules with stable stress-specific expression; here, we extend that approach to a cross-study, multi-stress classification context in a crop legume and showed that the SVM step contributed the most additional information precisely in the groups where consensus meta-analysis was most constrained by biological heterogeneity.

Panahi et al. (2024) applied Random Forest and C4.5 classifiers to 18 barley studies and identified 400 core stress-responsive genes using a precision-based discovery framework across a single stress type; our group-stratified design and inclusion of biotic stress conditions as separate groups instead produced substantially larger per-group core gene sets. Notably, classification precision was lowest for biotic root (0.198; permutation p = 0.02) – still significant, but well below the p < 0.001 threshold achieved in the other three groups. Class imbalance from a small positive-class training set is a recognized challenge that canaffect classifier performance in gene expression classification tasks (Al-Azani et al., 2024), and we attribute the reduced precision in this group to the small number of consensus meta-DEGs available as positive training labels combined with the mechanistic divergence of the three constituent studies. Thus, we interpreted results from this group with caution than those from the other three. More broadly, the 19 included studies span heterogeneous experimental designs and common bean genotypes, and applying a single set of consensus criteria across groups with fundamentally different heterogeneity profiles remains a constraint of our design; residual between-study heterogeneity likely contributed to the observed low biotic root consensus count.

### Stress tissue-group specificity and implications for molecular breeding

We did not find a single gene shared universally across all four stress-tissue groups, consistent with established stress specificity in plant transcriptome remodeling and indicating that a single unified common bean stress gene core does not exist based on this analysis. Abiotic root dominated the core gene count (4,178 genes, of which 3,420 were group-specific) with limited overlap with other groups, showing that root abiotic stress activates a largely independent transcriptional program relative to leaf abiotic or biotic responses. In the largest pairwise cross-group intersection we detected, where 38 upregulated and 13 downregulated genes are shared between abiotic root and biotic leaf, it represented only 1.9% and 3.0% of their respective upregulated gene sets, underscoring the degree of group independence. This pattern matches tissue- and stress-type-specific regulatory networks documented in model species including Arabidopsis (Sanchez-Munoz et al., 2025) and rice (Tarun et al., 2020) and is expected when stressors perceived by roots (soil-borne pathogens, drought, nutrient deficiency) activate distinct signaling cascades perceived by leaves (airborne pathogens, heat, light stress). We also observed an inverse relationship between consensus meta-DEG set size and ML-discovery proportion – highest in biotic root (80.1% ML-discovered) and lowest in abiotic leaf (73.2%), which further indicates that ML-based classification provides a non-redundant extension of meta-analysis results in high-heterogeneity groups rather than simply confirming the same genes. These patterns have direct implications for molecular breeding in that gene panels for abiotic root tolerance and biotic leaf resistance draw on largely non-overlapping transcriptional programs and likely need to be developed from separate expression data sources. Consequently, we would not expect genomic marker sets derived from abiotic root transcriptomics to hugely improve disease resistance prediction in leaf tissue or vice versa.

### Co-expression and protein-protein interaction networks define hub gene regulatory architecture across stress-tissue groups

Our WGCNA co-expression analysis revealed substantial differences in module structure across groups that reflect their underlying biological complexity. For example, we identified 14 co-expression modules in the biotic leaf group, with complete convergence between ML-identified and consensus hub genes across every module – indicating that the biotic leaf transcriptome is organized into multiple coordinated regulatory units, each independently captured by both discovery approaches. This high module count is consistent with the transcriptional complexity expected when studies representing bacterial blight, anthracnose, Fusarium wilt, and Rhizobium interactions each activate partially overlapping but pathogen-specific regulatory programs. The biotic root group, by contrast, was strongly ML-dominant: consensus meta-DEGs contributed hub genes to only one of three modules, further supporting our interpretation that biotic root defense gene expression is too variable across studies for consensus meta-analysis alone to recover the full hub gene landscape, and that ML-based classification fills this gap. We also found high co-expression overlap in abiotic leaf (mean 83.3 consensus meta-DEGs per ML hub gene module) alongside a consensus-exclusive module (brown; 9 consensus hub genes, 0 ML hub genes), together indicating that the abiotic leaf network combined a densely co-regulated core genes captured by both methods with a subset of regulatory genes recoverable only through meta-analysis consensus. ERF/AP2 domain transcription factors emerged as hub genes in three of four stress-tissue groups, consistent with the established role of ethylene response factors in integrating hormone signaling responses to both abiotic and biotic stressors across crop species. This aligns with the central role of ethylene signaling in multi-stress responses reported by Sanchez-Munoz et al. (2025) in Arabidopsis, although in our study ERF hub gene identification was based simply on cross-study meta-DEG and ML discovery rather than direct ethylene treatment.

The PPI analysis revealed divergent network architectures consistent with group-specific biology. Ribosomal proteins made up four of the five top hubs in the biotic leaf network, consistent with the role of translational reprogramming in plant immunity; this suggests that ribosomal subunit composition changes represent a mechanism by which the biotic leaf stress response modulates defense-related protein production (Bach-Pages et al., 2025). In the abiotic root network, we found that salicylic acid signaling (NPR1-like, Phvul.006G131400) and jasmonate perception (COI1, Phvul.008G226500) occupied the highest-connectivity positions, indicating that biotic-defense hormone pathway components are co-opted into the abiotic root stress response. This is a cross-stress signal integration pattern consistent with documented SA-JA pathway activation under drought and heavy metal stress in legumes (Mohi-Ud-Din et al., 2021; Pandey et al., 2025). The ML-discovered chaperone Hsp90 (Phvul.004G107700) also ranked among the top abiotic root PPI hubs, consistent with its function in stabilizing NLR immune receptors and client proteins under drought conditions (Madirov et al., 2025; Yang et al., 2026). Hub genes in the biotic root group were predominantly hormone signaling genes (39.8%), matching the expected pattern of jasmonate-ethylene pathway activation in root defense against necrotrophic pathogens and symbiotic microbes. Notably, no gene was identified as a PPI hub in more than one stress-tissue group, indicating that the network interaction landscape is as group-specific as the transcriptional landscape itself. We also detected disease resistance (R) genes in the abiotic leaf hub gene set (n = 2), consistent with documented cross-talk between abiotic and biotic stress defense pathways in legumes.

Functional enrichment analysis identified significant GO and KEGG enrichment only in the abiotic leaf hub gene set, with peroxidase activity independently confirmed by both GO (GO:0004601, GO:0006979) and KEGG (K00430) analyses; ROS scavenging via peroxidases is a conserved abiotic stress tolerance mechanism across plant species (Pandey et al., 2017), and the enrichment of WRKY transcription factors fits their established role in regulating abiotic stress-responsive gene expression (Kiranmai et al., 2018). We interpret the absence of significant enrichment in the other three groups as reflecting the functional heterogeneity of hub genes identified across diverse biotic stressors combined with smaller hub gene counts in those groups, rather than a true absence of coordinated function. Also, the statistical threshold stringency as applied here for declaring enrichment may be worth consideration especially for biological context.

### Cross-species ortholog conservation supports functional transfer of stress-responsive gene evidence to cowpea

We identified *V. unguiculata* counterparts for 238 of 291 ML hub genes (81.8%) across the abiotic and biotic stress groups, with mean sequence identity for Tier 2 orthologs of 87.0-88.0%, indicating high protein-coding conservation between the two species at the hub gene level. Functional class proportions were substantially conserved between *P. vulgaris* hub genes and their cowpea orthologs: Kinase Signaling (12.6%), Transcription Factor (10.9%), and Transporter (6.7%) classes were all represented in cowpea at proportions comparable to common bean, and the proportional distribution across non-other functional classes was structurally similar between species. We interpret this degree of cross-species functional conservation as evidence that the transcriptional regulatory architecture underlying stress responses is broadly shared between these two legume species, consistent with the extensive genomic synteny documented between common bean and cowpea (Lonardi et al., 2017), providing biological justification for using common bean meta-analysis results to prioritize cowpea markers. This supports our hypothesis that gene-proximal SNP subset selection informed by common bean expression data carries information relevant to cowpea phenotypic prediction. Notably, the Other/Unknown class constituted 69.3% of cowpea hub gene orthologs (165 of 238), reflecting the narrow scope of the functional classification scheme rather than a genuine lack of functional annotation as cowpea’s genome annotation resource is almost complete but many annotated genes do not fall within the specific functional categories examined here. Thus, we infer functional equivalence between common bean and cowpea orthologs from sequence homology and class assignment, but this has not been experimentally verified at the protein interaction or promoter regulation level.

### External validation confirms transferability of ML-predicted drought-responsive gene sets

Here, we explored if a drought-specific classifier can be useful for identifying drought-responsive genes from dataset of any common bean drought-stress study without training with the dataset. Essentially, we validated core drought gene set against an independent common bean drought experiment and confirmed that it is transferable beyond the training studies (Odds Ratio = 1.35; p = 0.001), with higher predicted drought-response probabilities in external DEGs than in non-significant genes. The enrichment odds ratio, while statistically significant, reflects a moderate effect size, which we attribute to experimental design differences between the training studies (treatment, diverse platforms and genotypes) and the external experiment (genotype, two time points, single greenhouse environment). Consequently, we would expect these design differences to reduce the transferability of any expression-based model trained on multi-study meta-features. The AUC (0.967), by contrast, indicates near-complete separation of external DEGs from non-DEGs by predicted drought-response probability rankings, showing that the probability ranking remains broadly informative for identifying drought-responsive genes even when the experimental context differs from the training data. However, we note that this estimate comes from a single external experiment, so we cannot rule out that some of the separation reflects dataset-specific differences between external DEG and non-DEG sets rather than generalizable drought-response signal. Essentially, testing the gene set from the prediction model against additional, multiple independent drought experiments would help confirm the robustness of the model’s performance. Meanwhile, leave-one-study-out cross-validation (LOSO-CV) across the four studies used for the drought-specific classifier further supported transferability (AUC ranged from 0.73 - 0.98).

### RNA-seq-informed SNP subset selection improved genomic prediction accuracy for multiple stress traits in common bean and cowpea

As agricultural systems face increasing pressure from climate change and emerging pathogen variability, the ability to integrate multi-study multi-modal directly into molecular breeding tools has direct implications for crop improvement programs in legumes like common bean and cowpea, two protein-rich legumes of global food security relevance. Here, our genomic prediction analysis provided direct experimental evidence that RNA-seq-derived gene lists identified herein carries marker-level information useful for phenotypic prediction in two legume species.

In common bean, the Meta-DEG SNP subset consistently ranked highest for SCN resistance and bacterial wilt Race 528 and Race 557, though absolute improvements over baseline were small (0.5%-3.3%). We attribute this to SCN prediction accuracy already being high at baseline (r = 0.759-0.831), leaving limited room for improvement once the genomic relationship matrix already captured most of the additive genetic variance with the full SNP set. Our random draw comparison confirmed that Meta-DEG-proximal SNPs exceeded the null distribution for Race 528 (r = 0.551 vs. random mean r = 0.546) and Race 557 (r = 0.456 vs. random mean r = 0.440), but not Race 597 (r = 0.446 vs. random mean r = 0.451) – indicating that the gene expression-derived evidence set is informative for races whose resistance genetics overlap with the stress transcriptome, and uninformative where resistance is instead governed by regions/loci outside the stress-responsive gene landscape. We suspect the underperformance of all evidence-based schemes for Race 597 reflects either a small number of major-effect resistance loci not enriched in the stress transcriptome, or that the expression studies informing our SNP subset assignments were not representative of the genetic architecture underlying resistance to that race. Transferring common bean SNP subset assignments from gene expression data to disease resistance prediction – two biologically distinct processes, remains an inherent limitation of our cross-phenotype approach; nonetheless, the overall pattern of evidence-based schemes ranking above Background across most traits supports our hypothesis that common bean stress-responsive genes harbor SNPs with functional relevance beyond their transcriptomic context.

In cowpea, the ML gene-proximal SNP subset achieved the highest prediction accuracy for five of six traits, with the largest gains observed for traits with the lowest baseline accuracy, RTI_H: r = 0.104 vs. r = 0.056 for baseline GBLUP (+86.7%), indicating that gene-set-based subset selection is most impactful when baseline GBLUP captures little of the genetic variance, likely due to small sample size or low trait heritability and/or environmental effect. For traits with moderate baseline accuracy (RTI_C: r = 0.438; RTI_C_T: r = 0.449; RTI_C_U: r = 0.516), ML gene improvements were smaller (0.9%-3.1%), matching our expectation that prediction gains from SNP subset selection diminish as the baseline genomic relationship matrix captures an increasing share of the additive genetic variance. Notably, for the composite drought tolerance score (dt_Score), the Meta-DEG SNP subset outperformed the ML gene subset (+5.4% vs. −1.4%) – the only trait where this ranking reversed, suggesting that the broader consensus gene set better captured the polygenic architecture of composite tolerance, while the more refined ML hub gene set may include SNPs in functionally specific genes less relevant to a composite score, diluting its signal. Our random SNP draw comparison confirmed that ML gene-proximal SNPs exceeded the null distribution in five of six traits, while Meta-DEG-proximal SNPs exceeded random in only two of six (dt_RTI_C_T and dt_Score), establishing that the accuracy advantage of ML gene-proximal SNPs reflects the biological informativeness of the gene set rather than a statistical artifact of using fewer or random SNPs.

While we are able to show the biological significance of the transcriptome evidence used in this genomic prediction design, we would like to highlight some other limitations. For example, we conducted the bacterial wilt analysis using the lower-density 6K BARCBean6K3 SNP panel and we evaluated SCN resistance and cowpea abiotic traits using a moderate-density panels (87,176 and 103,916 SNPs, respectively) based on available dataset and to reflect what was reported on the two disease traits (Zia et al. 2022; Shi et al. 2025). Consequently, the BW panel may not offer sufficient resolution to capture the full benefit of gene-proximal SNP subset selection, and we would expect higher-density genotyping data to produce larger accuracy gains for that trait system. We also used a single GRM kernel per evidence tier throughout; multi-kernel approaches combining evidence and background GRMs within the same model such as genomic feature BLUP (Edwards et al., 2016; Ye at al. 2020), may capture a larger proportion of genetic variance and this warrant evaluation in future work.

## Conclusion

In summary, we showed that RNA-seq meta-analysis, machine learning, and network analysis can be integrated into a unified pipeline for identifying stress-responsive gene sets in common bean, and that these gene sets carry marker-level information relevant to genomic prediction. Our group-stratified design, which separated abiotic and biotic conditions across leaf and root tissues, revealed that stress-tissue specificity is a consistent feature of the common bean stress transcriptome, and that ML-based gene discovery provides the most complementary information precisely in groups where consensus meta-analysis is most constrained by biological heterogeneity. We expect the approach presented here to be applicable to other crop species for which multi-study RNA-seq data are available in public databases, and our cross-species knowledge transfer component extends the utility of single-species transcriptomics beyond the source species. Looking forward, higher-density genotyping data, multi-kernel GRM designs, and tissue-specific expression panels matched to the target trait system may further improve the genomic prediction gains we demonstrated here.

## Method

### Study selection and data retrieval

RNA-seq datasets were retrieved from the National Center for Biotechnology Information Sequence Read Archive (NCBI SRA) (https://www.ncbi.nlm.nih.gov/bioproject/) . A structured keyword search was performed using the following query: ("drought"[All Fields] OR "water deficit"[All Fields] OR "heat"[All Fields] OR "cold"[All Fields] OR "salt"[All Fields] OR "nacl"[All Fields]OR "osmotic"[All Fields] OR "heavy metal"[All Fields] OR "bacterial"[All Fields] OR "fungal"[All Fields]) OR "virus"[All Fields] OR "microbes"[All Fields] AND "Phaseolus vulgaris"[porgn] AND "Expression profiling by high throughput sequencing"[Filter]. This search resulted in a total of 22 independent studies of which 19 was retained, encompassing a diverse range of abiotic and biotic stress types. Details of the selected data sets are presented in Table S1. Studies were retained if they: (i) used *P. vulgaris* as the experimental organism; (ii) included stress treatment and corresponding non-stressed control; (iii) provided a minimum of two biological replicates per condition; and (iv) covered leaf or root tissue or a combination of both. Manual curation of database metadata confirmed stress treatment conditions, sample tissue identity, and pairing of treated and control samples within the same project. FASTQ files for each SRA run accession were retrieved using the prefetch utility of the SRA Toolkit (v3.0.0). (https://github.com/ncbi/sra-tools). Essentially, the final list was composed of 461 transcriptome datasets from 19 studies.

### Transcript quantification and differentially expressed gene (DEG) detection

Raw read quality was assessed with FastQC (v0.12.1) and summarized across studies with MultiQC (v1.21). Transcript quantification was performed using Salmon (v1.10.0; Patro et al. 2017) against the *P. vulgaris* v2.1 reference transcriptome (Schmutz et al. 2014). The reference genome was selected based on annotation completeness and continuous curation since assembly. Transcript-level abundance estimates were summarized to gene-level counts using the tximport R package (Soneson et al. 2015), with counts derived from transcripts per million (TPM) scaled by effective length. Batch effects within studies were corrected using ComBat-seq prior to differential expression testing (Leek et al., 2012). To preserve the biological context within each study, DESeq2 models (Love et al. 2014) were fitted according to the study-specific experimental design of each dataset. Genes with counts ≥ 10 in fewer than three samples were removed and differentially expressed genes were defined by a false discovery rate (FDR) cut-off of < 0.05 and absolute log2 fold change (|LFC|) ≥ 1.5. Principal component analysis and Spearman sample-correlation heatmaps were produced pre- and post-batch correction for quality control.

### Cross-study hierarchical clustering analysis

Cross-study gene overlap was quantified by constructing gene-presence matrices for up- and down-regulated DEGs and visualized as heatmaps using pheatmap R package (Kolde, 2019). Gower dissimilarity (Gower, 1971) was computed on a study-by-gene binary matrix and used to perform agglomerative hierarchical clustering with both complete and average linkage methods. Also, Baker’s gamma was computed to assess dendrogram concordance between the linkage methods (Galili 2015). Finally, a diverging horizontal barplot was used to summarize per-study DEG counts.

### Stress-tissue group definition, meta-analysis approach and consensus gene designation

Each study was assigned a stress-tissue group identifier to account for the distinct transcriptomic signatures associated with stress type and tissue of origin. The identifier was defined as the combination of stress category (abiotic or biotic) and tissue type (leaf or root), yielding four groups (abiotic_leaf, abiotic_root, biotic_leaf, and biotic_root). Next, full results including log2FoldChange, lfcSE, and unadjusted p-values, were extracted per study and pooled by stress-tissue group. Essentially, per-study results were retained as independent observations at the gene level, with each gene’s effect estimate reflecting the contrast fitted under that study’s experimental design. Only genes represented in at least three studies within a group were included to ensure that meta-analytic estimates were based on independently replicated evidence. Meta-analysis was performed independently for each stress-tissue group using a dual-criterion approach. First, Fisher’s combined probability statistic was computed *x*^2^ = −2 ∑*log*(*p_i_*), distributed as chi-squared with df = 2k, where k is the number of contributing studies and *p_i_* are per-study unadjusted p-values. Second, a random-effects model was fitted by Restricted Maximum Likelihood (REML) implemented in the R package, metafor (Viechtbauer, 2010), with per-study log2FoldChange as effect sizes and per-study lfcSE squared as sampling variances. The random-effects framework was used in addition to the Fisher’s because studies differed in genotype, growth conditions, and stress severity, violating the assumption of a common true effect size. Between-study heterogeneity was quantified by Cochran’s Q statistic, I² index, and the between-study variance (τ²). Benjamini-Hochberg FDR correction was applied to Fisher and REML meta p-values separately within each group.

Consensus meta-DEG significance designation was assigned by a set of I²-stratified rules to balance sensitivity and specificity across the heterogeneity spectrum: genes with low heterogeneity (I² < 50%) required Fisher FDR < 0.05 and REML FDR < 0.05; genes with moderate heterogeneity (50% ≤ I² < 75%) required Fisher FDR < 0.05 and REML p < 0.10 (Relaxed criterion, as moderate heterogeneity can inflate the REML standard error and cause genuinely reproducible effects to fail the stringent FDR threshold); and genes with high heterogeneity (I² ≥ 75%) required Fisher FDR < 0.01 and directional consistency ≥ 0.80 across contributing studies (proportion of studies with expression in the majority direction), since high heterogeneity renders pooled effect size unreliable but supermajority directional agreement across studies could constitutes strong reproducibility evidence. Genes for which REML could not be fitted were excluded from the consensus.

To confirm that consensus (meta-DEG) calls exceeded what would be expected by chance under the null hypothesis of no cross-study association, the empirical significance of observed consensus gene counts was assessed by permutation; the log_2_FoldChange and unadjusted p-values were independently permuted 1,000 times simultaneously across all genes and studies, the full dual meta-analysis was re-run under the permuted data, and the number of consensus-significant genes was recorded. Finally, an empirical p-value was computed as the proportion of permutations in which the permuted count equaled or exceeded the observed count.

### Machine learning gene discovery and validation

Two machine learning (ML) analyses were implemented. For each stress-group, the feature matrix comprised three variables derived from the group-specific meta-log_2_FoldChange: the effect estimate, its square, and its absolute value; a parsimonious representation was used because the group-stratified meta-analysis produces a single effect estimate per gene per group, precluding richer feature construction at this stage. A Support Vector Machine (SVM) classifier (using R package, e1071) with radial basis kernel was fitted using nested cross-validation: within each outer fold, inner hyperparameter tuning was performed by three-fold cross-validation over cost values (1, 10) and gamma values (0.01, 0.1, 1). An outer five-fold cross-validation loop provided unbiased estimates of precision. The classification precision was computed as TP/(TP + FP) = consensus/( consensus + ML-discovered). Also, accuracy, recall, F1, specificity, and area under the receiver operating characteristic curve (AUC) were reported. Genes with consensus meta-DEG status in any of the four groups were labeled as pre-classified (class 1); all remaining genes were labeled class 0. The proportion of class 1 genes in the combined gene set (0.169) served as the theoretical precision baseline, representing the expected precision of a classifier that assigns positive labels at random. SVM output was used to identify class transitions: 1 to 1 (confirmed) genes are pre-classified genes retained as class 1 by SVM, and 0 to 1 (ML-discovered) genes are those reclassified from 0 to 1 by SVM. Class imbalance was addressed by assigning a fixed weight of 3 to the minority positive class relative to 1 for the majority negative class, to prevent the classifier from defaulting to the majority negative class. Each gene was assigned to exactly one outer test fold; genes predicted as stress-responsive by the SVM in their assigned fold that were absent from the meta-analysis consensus (0 to 1 transitions) were designated ML-discovered genes, representing biologically relevant candidates below the conventional significance threshold. Genes confirmed by both meta-analysis and SVM (1 to 1 transitions) were designated ML-confirmed genes.

To assess whether genes identified by a drought-specific machine learning classifier overlapped with differentially expressed genes from our independent common bean drought experiment, a drought-specific classifier was trained on genes drawn from five drought studies (three leaf and two root). The feature matrix comprised nine variables: meta-log_2_FoldChange from the combined, leaf-specific, and root-specific meta-analyses; the absolute meta-log_2_FoldChange; the number of studies in which the gene was observed; I²; between-study standard deviation of log_2_FoldChange; direction consistency; and a range-normalized Fisher combined statistic. Features were standardized to zero mean and unit variance prior to model fitting. A Random Forest classifier (ranger) was fitted as the primary model using five-fold cross-validation and a support vector machine was fitted (like previously) in parallel as a secondary classifier. Class imbalance was addressed by downsampling the majority class within each cross-validation fold and setting the positive-class weight to the ratio of negative to positive class observations, capped at a maximum of 10. Core drought-responsive genes were defined as all genes predicted positive in their held-out cross-validation fold by the Random Forest, irrespective of pre-classification status. Generalization was assessed by leave-one-study-out cross-validation (LOSO-CV), in which each of the five drought studies was held out in turn as the test set, with the Random Forest retrained on the remaining studies using three-fold cross-validation. Permutation testing (1,000 permutations of class labels) established an empirical p-value for the observed five-fold cross-validated F1 score, confirming that classifier performance exceeded chance expectation. External validation was performed using DEGs from our independent RNA-seq drought study, by a one-sided Fisher’s exact test of core gene enrichment among those externally detected DEGs, and by ROC-AUC of Random Forest probability scores against their differential expression status.

### Weighted gene co-expression network analysis (WGCNA)

Weighted gene co-expression networks were constructed independently for each of the four stress-tissue groups using consensus-significant genes as input, with per-study log_2_FoldChange values as the expression matrix and missing values set to zero (Zhang and Horvath, 2005). Soft-thresholding power was selected based on the scale-free topology criterion (R² > 0.80). Signed topological overlap matrices (TOM) were computed and subjected to hierarchical clustering with average linkage. Modules were identified by dynamic tree cutting and modules with eigengene dissimilarity < 0.25 were merged. Hub genes within each module were ranked by intramodular connectivity. To validate ML-discovered genes (0 to 1 transitions), each ML-discovered gene was projected onto the consensus-anchored network by correlating its expression profile with module eigengenes and assigned to the module with the highest absolute correlation. ML-discovered genes ranking in the top 10 by intramodular connectivity within their assigned module and with module eigengene correlation |r| > 0.80 were designated priority ML candidates, providing network-level validation independent of differential expression testing.

### Protein-Protein interaction network analysis

PPI networks were constructed for each stress-tissue group using the STRINGdb R package (version 12.0). WGCNA hub genes from each group were mapped to STRING protein identifiers and their subnetworks extracted. Degree centrality was computed using the igraph R package, and the top 12 genes per group by degree were designated PPI network hubs. Hub gene meta-log_2_FCprofiles were visualized as a heatmap across the four stress-tissue groups.

### Functional annotation of genes

Hub genes were annotated using the *P. vulgaris* v2.1 gene annotation (Phytozome v13; Pvulgaris_442_v2.1.P14), with Pfam domain assignments (Pfam database, accessed January 2026), KEGG Orthology (KO) terms, and Panther superfamily classifications. Annotated genes were classified into broad functional gene family categories. Enrichment of consensus meta-DEGs against the P. vulgaris annotated gene background was tested for each stress-tissue group by one-sided Fisher’s exact test, applied separately to GO terms, KO terms, Pfam domains, and Panther superfamilies. Enrichment p-values were FDR-adjusted by the Benjamini–Hochberg method and terms with FDR < 0.05 were retained.

### Independent drought experiment

#### Plant materials and growth conditions

Data from an independent common bean drought experiment previously conducted at the Harry R. Rosen Alternative Pest Control Centre, University of Arkansas, Fayetteville, AR, were used to evaluate the external validity of the drought-specific gene set. The experiment consisted of two independent greenhouse trials carried out under average conditions of 61.9% relative humidity and 27℃ air temperature. The common bean genotypes used in this study were selected based on performance in a previous study which had evaluated a set of common bean accessions from the USDA core collection. PI549793 (originally from China) and PI361408 (originally from India) showed moderate tolerance and complete susceptibility respectively in the mini-core set evaluated in the studies (Yang et al. 2019). For the trials in this study, direct seeding was done in 7.5 cm spaced rows within soil mix-filled polypropylene boxes (Sterilite Corporation, Townsend, MA) with dimensions 88.6 × 42.2 × 15.6 cm. For the first trial, each genotype was directly seeded into two rows at six plants/row and assigned rows randomly across the trial. In the second trial, each genotype was directly seeded into five rows at four plants/row, assigned all rows randomly within the box. To introduce moisture stress, watering was stopped in 19-days-old seedlings, while watering was continued for the rest of the seedlings. The volumetric water content of the soil mix was then monitored using a soil moisture meter (VG-METER-200, Vegetronix) for both water regimes each day. After 2 days following water withdrawal, we started collection of data on drought-associated traits including total chlorophyll value, leaf relative water content, days to flowering, days to pod emergence, number of dead plants and recovery rate.

#### Physiological measurements

For chlorophyll content, total chlorophyll value was assessed using soil plant analysis development (SPAD-502) Plus meter (Spectrum Technologies, Inc., Plainfield, IL) on 3rd trifoliate leaves of each plant for 10 consecutive days. Measurements were taken approximately same time on each day. For each plant, each trifoliate leaves were assessed for chlorophyll value at three different points and the average value was recorded. To assess relative water content in the leaves of both genotypes over 10 days under water stress, we carefully collected four leaves from four separate plants of each genotype. Leaves were collected on the 1st, 3rd, 6th, 8th, and 10th day after introduction of moisture stress. Each leaf was immediately placed inside a Ziplock plastic bag and kept in a cool container until used for fresh weight measurement followed by immersion in distilled water inside beaker for 24 hours under room temperature. Turgid weight of individual leaf was measured, and leaves were placed in a laboratory oven overnight at 65℃. Overnight dried leaves were used for dry weight measurement. Phenotypic data from total chlorophyll value and relative water content were analyzed using the JMP software (SAS Institute). The chlorophyll response was analyzed as a univariate analysis of variance (ANOVA) using the Split-Plot repeated measures approach within the JMP software (SAS Institute). Water regime was considered whole plot effect while genotype effect was nested within water regime and the term specified as a random effect. The restricted maximum likelihood method (REML) was adopted for fitting the linear mixed model. Other measurements collected including days to flowering and pod emergence was collected 20 days after moisture stress introduction.

#### Sample collection for transcriptional analyses

We collected leaf samples from the contrasting genotypes during the seedling stage at three different time points (24, 144 and 192 hours after drought treatment) for transcript profiling. Leaf samples were carefully collected from each of the genotypes throughout the different time point. We collected at least five leaf (bulked) samples from plants of each genotype at each time point, where each bulked leaf samples represented a replicate.

#### RNA-seq library preparation and sequencing

A total amount of 1 µg RNA per sample was used as input material for the RNA sample preparations. Sequencing libraries were generated using NEBNext® UltraTM RNA Library Prep Kit for Illumina® (NEB, USA) following manufacturer’s recommendations and index codes were added to attribute sequences to each sample. Briefly, mRNA was purified from total RNA using poly-T oligo-attached magnetic beads. Fragmentation was carried out using divalent cations under elevated temperature in NEBNext First Strand Synthesis Reaction Buffer (5X). The first strand cDNA was synthesized using random hexamer primer and M-MuLV Reverse Transcriptase (RNase H-). Second strand cDNA synthesis was subsequently performed using DNA Polymerase I and RNase H. The remaining overhangs were converted into blunt ends via exonuclease/polymerase activities. After the adenylation of 3’ ends of DNA fragments, NEBNext Adaptor with hairpin loop structure were ligated to prepare for hybridization. To select cDNA fragments of preferentially 150–200 bp in length, the library fragments were purified with AMPure XP system (Beckman Coulter, Beverly, USA). Then, 3 µl USER Enzyme (NEB, USA) was used with size-selected, adaptor-ligated cDNA at 37°C for 15 min followed by 5 min at 95°C before PCR. PCR was performed with Phusion High-Fidelity DNA Polymerase, Universal PCR primers and Index Primer. Finally, PCR products were purified (AMPure XP system) and library quality was assessed on the Agilent Bioanalyzer 2100 system. Libraries were clustered on a cBot Cluster Generation System using PE Cluster Kit cBot-HS (Illumina) and sequenced on an Illumina platform to generate paired-end reads.

#### Bioinformatic analysis

Transcript-level quantification was performed using Salmon and imported to gene-level counts using the tximport R package, with transcript-to-gene mapping derived from the P. vulgaris v2.1 reference transcriptome. Results from differential expression analysis on timepoints were reported but only T3 (144 hours) and T4 (192 hours) were used for validation. Genes with total read counts below 10 across all samples were removed. Differential expression was performed using DESeq2 like other studies. Genes were declared differentially expressed at a Benjamini-Hochberg adjusted p-value < 0.05 and |log_2_FC| ≥ 2.

KO and GO term mappings were obtained from the P. vulgaris v2.1 gene annotation file (Pvulgaris_442_v2.1.P14.annotation_info.txt; Phytozome v14). KEGG pathway enrichment was performed using the enrichKEGG function in the clusterProfiler R package (p-value cutoff = 0.05) on four gene sets: sustained drought-responsive genes, tolerant genotype-specific genes, susceptible genotype-specific genes, and shared genes expressed in both genotypes. GO term enrichment for sustained drought-responsive genes was performed using the enricher function with a custom TERM2GENE table constructed from GO annotations in the same Phytozome file (*p*-value cutoff = 0.05). For tolerant-specific and susceptible-specific gene sets, GO annotations were additionally retrieved from Ensembl Plants via the biomaRt R package and GO enrichment was repeated.

### Ortholog mapping and gene tier stratification

Consensus meta-DEGs and ML-discovered genes from *P. vulgaris* were mapped to orthologous genes in *Vigna unguiculata* via Ensembl Plants BioMart, retaining one-to-one ortholog relationships only. Next, genes were stratified into four tiers: Tier 1, meta-analysis consensus DEGs; Tier 2, WGCNA hub genes from ML-discovered gene sets; Tier 3, remaining ML-discovered genes; Tier 4, all other reference genome genes. Abiotic and biotic tiers were constructed independently by combining leaf and root gene lists. Results from the ortholog mapping are presented.

### Validation of genes via genomic prediction in common bean and cowpea

Genomic best linear unbiased prediction (GBLUP) was used to evaluate whether SNP subsets defined by transcriptomic evidence from the meta-analysis and ML frameworks, assigned to tiers based on proximity, yielded higher prediction accuracy than equal-sized random SNP subsets drawn from the same panel, thereby isolating the biological contribution of the gene annotation from the effect of marker density alone. Genomic coordinates for tier-assigned genes were retrieved from Ensembl Plants, and 50-kb windows were defined around each gene body; SNPs falling within one or more windows were assigned to the highest-confidence tier among overlapping windows, and the same window-based assignment was repeated using cowpea ortholog genomic coordinates for cross-species validation. Four SNP subsets were evaluated in all analyses: Meta_DEG (Tier 1 SNPs proximal to consensus meta-DEG gene bodies), ML_genes (Tier 2 and Tier 3 SNPs proximal to ML-discovered gene bodies), Background (SNPs outside all evidence gene windows), and Unweighted (all SNPs; standard GBLUP baseline), each used to construct a separate genomic relationship matrix (GRM) by the VanRaden (2008) method. In all analyses, a single-trait GBLUP model was fitted as thus:

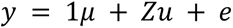

where y is the phenotypic vector, μ is the overall mean, u ∼ N(0, Gσ²g) is the vector of genomic breeding values, and e ∼ N(0, Iσ²e) is the residual vector.

For common bean, two independent datasets were used: a soybean cyst nematode (SCN) resistance study (association analysis including genomic prediction) on 354 USDA accessions with 87,176 SNPs derived from whole genome resequencing, and with phenotypic records for three SCN HG types (HG_Type_7, HG_Type_2.5.7, and HG_Type_1.3.6.7) (Shi et al. 2025), and a bacterial wilt resistance study (association analysis including genomic prediction) on 168 accessions genotyped with the BARCBean6K3 SNP panel (approximately 6,000 SNPs), with phenotypic records for three pathogen races (BW_528, BW_557, and BW_597) (Zia et al. 2022). For the bacterial wilt dataset, SNP matrices were filtered by excluding markers with missing rate exceeding 20%, heterozygosity rate exceeding 10%, or minor allele frequency below 0.05. Genomic estimated breeding values (GEBVs) were obtained by REML implemented in the sommer R package. Prediction accuracy was computed as the Pearson correlation (r) between observed phenotypic values and GEBVs in the held-out validation set.

For cowpea, the SNP panel comprised 110,000 markers derived from whole genome resequencing and uniformly subsampled at 10,000 SNPs per chromosome across all 11 cowpea chromosomes for 253 USDA accessions (Ravelombola et al. 2021; 2025); markers on unplaced scaffolds were excluded, and sequential quality control removed markers with missing rate exceeding 20%, heterozygosity exceeding 10%, or minor allele frequency below 0.05, retaining 103,916 SNPs. Six abiotic stress phenotypes were analyzed, expressed as relative tolerance indices (RTI = 100 × stressed value / non-stressed value): fresh biomass (st_RTI_FB), unifoliate leaf SPAD chlorophyll (st_RTI_C), and plant height (st_RTI_H) under salt stress; and unifoliate leaf SPAD chlorophyll (dt_RTI_C_U), first trifoliate leaf SPAD chlorophyll (dt_RTI_C_T), and a composite drought tolerance score (dt_Score). GRMs were similarly constructed using the VanRaden (2008) method.

Prediction accuracy was estimated by five-fold cross-validation repeated 100 times, with accuracy per repeat computed as the mean Pearson r across folds and final accuracy reported as the mean ± SE across 100 repeats; model calibration was assessed identically to common bean. To establish size-matched null distributions, 100 random GRMs were constructed at the Meta_DEG SNP count and the ML_genes SNP count (drawn without replacement) and each evaluated under the same five-fold CV and 100-repeat procedure.

## Acknowledgments

The authors sincerely thank all scientists and collaborators who contributed to this project, as well as the reviewers and editors for their constructive feedback. AI-based coding assistants were used during the development and debugging of analysis scripts; all analytical decisions, script execution and interpretation of results were performed and verified by the authors.

## Author contributions

D.O. and A.S. conceptualized and designed the project. D.O., L.R., O.A., B.K. performed data mining and transcriptome quantification. D.O. conducted phenotyping for independent drought study, performed all other analyses and wrote the manuscript with revisions from all authors. S.K. reviewed and provided helpful insight into initial results. Y.Y. contributed to an initial drought study experiment. W.R. contributed the cowpea datasets for genomic prediction. A.S. supervised the project and secured funding.

## Data availability

The datasets supporting the conclusions of this article are available in the NCBI Sequence Read Archive (SRA) repository under the corresponding BioProject ID of individual study. The scripts used for the analysis can be found at https://github.com/dotunolaoye/CB_RNA_Meta_Analysis_project/. An interactive Shiny dashboard for exploring the meta-analysis and genomic prediction results is available at https://dotunolaoye.shinyapps.io/CB_RNAseq_dashboard/

---

**Table S1.**
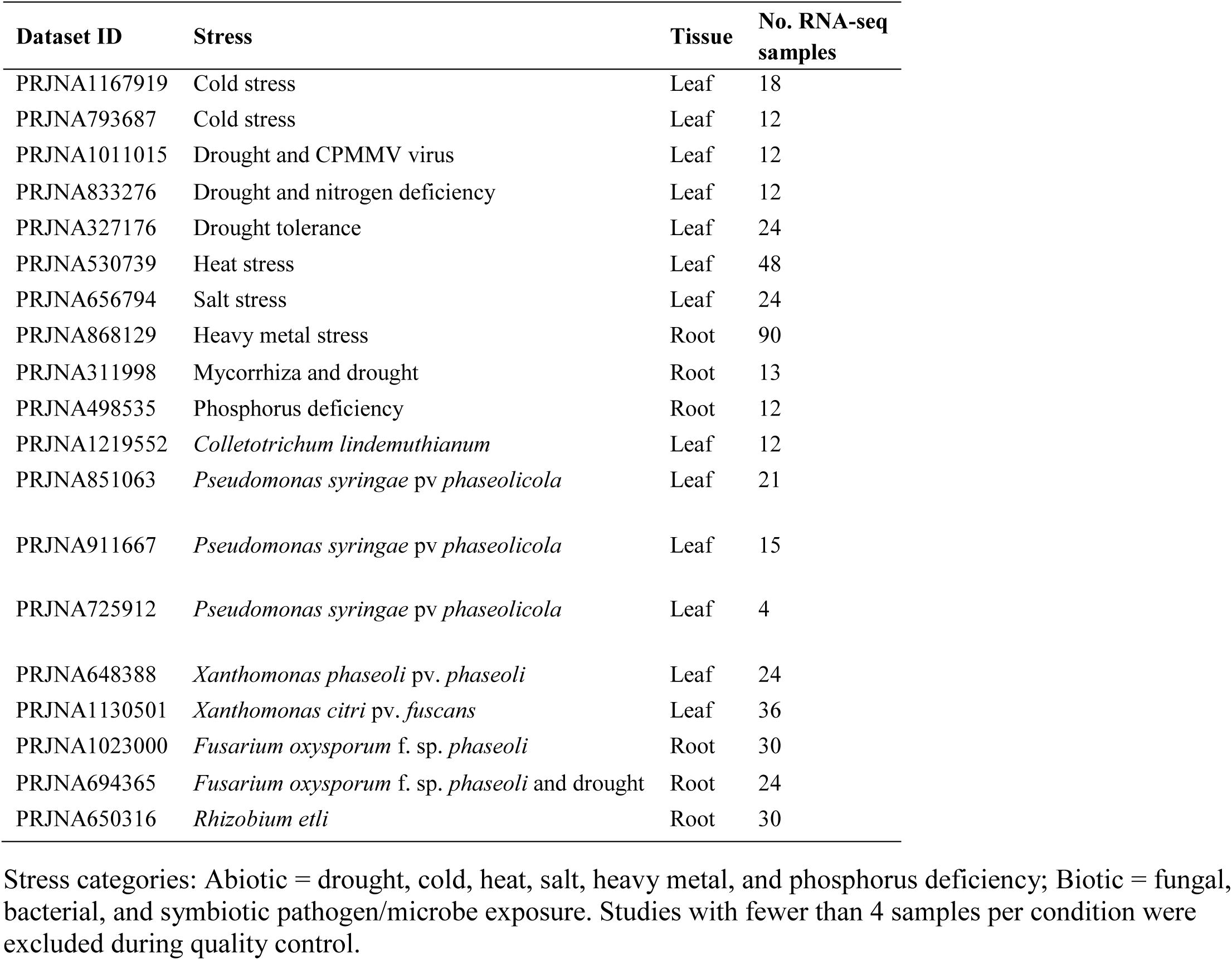
Characteristics of the 19 RNA-seq studies included in the meta-analysis. This summarize the abiotic and biotic stress conditions to captured in the analysis.

**Table S2.** Leave-one-study-out cross-validation (LOSO-CV) performance of the Random Forest drought-response classifier across the four studies.

| Held-out study | Dataset ID | Tissue | Test DEGs | AUC |
| --- | --- | --- | --- | --- |
| Drought tolerance | PRJNA327176 | Leaf | 3,695 | 0.989 |
| Drought and nitrogen deficiency | PRJNA833276 | Leaf | 3,739 | 0.981 |
| Drought and CPMMV virus | PRJNA1011015 | Leaf | 10,233 | 0.848 |
| Fusarium wilt and drought | PRJNA694365 | Root | 127 | 0.735 |

Test DEGs = number of significant differentially expressed genes in the held-out study; AUC = area under the ROC curve for classification of held-out-study genes.

**Table S3.** Descriptive statistics for SPAD-measured chlorophyll values in *P. vulgaris* genotypes under drought stress and well-watered conditions in two greenhouse trials.

|  | Treatment<sup>a</sup> | Mean | Standard Error | Median | Standard Deviation | Kurtosis | Skewness | Range |
| --- | --- | --- | --- | --- | --- | --- | --- | --- |
| PI361408 | Trial1_D | 23.0 | 0.9 | 24.3 | 6.0 | -0.9 | -0.4 | 22.9 |
|  | Trial1_W | 29.4 | 0.3 | 29.5 | 2.2 | 2.4 | -1.1 | 10.7 |
|  | Trial2_D | 33.6 | 0.2 | 33.6 | 1.8 | -0.3 | 0.1 | 8.3 |
|  | Trial2_W | 39.6 | 0.1 | 39.4 | 1.5 | -0.9 | 0.3 | 4.9 |
| PI549793 | Trial1_D | 28.1 | 0.4 | 29.0 | 2.8 | -1.3 | -0.4 | 8.6 |
|  | Trial1_W | 31.8 | 0.4 | 31.8 | 2.3 | -0.6 | 0.0 | 9.2 |
|  | Trial2_D | 37.3 | 0.2 | 36.9 | 2.2 | -1.0 | 0.3 | 7.8 |
|  | Trial2_W | 39.7 | 0.1 | 39.2 | 1.2 | -1.3 | 0.6 | 3.4 |

<sup>a</sup>D: Drought or water-deficit condition; W: Well-watered condition
The values for the first trial are from two replicates and the values for the second trial are from four replicates.

**Figure S1.**
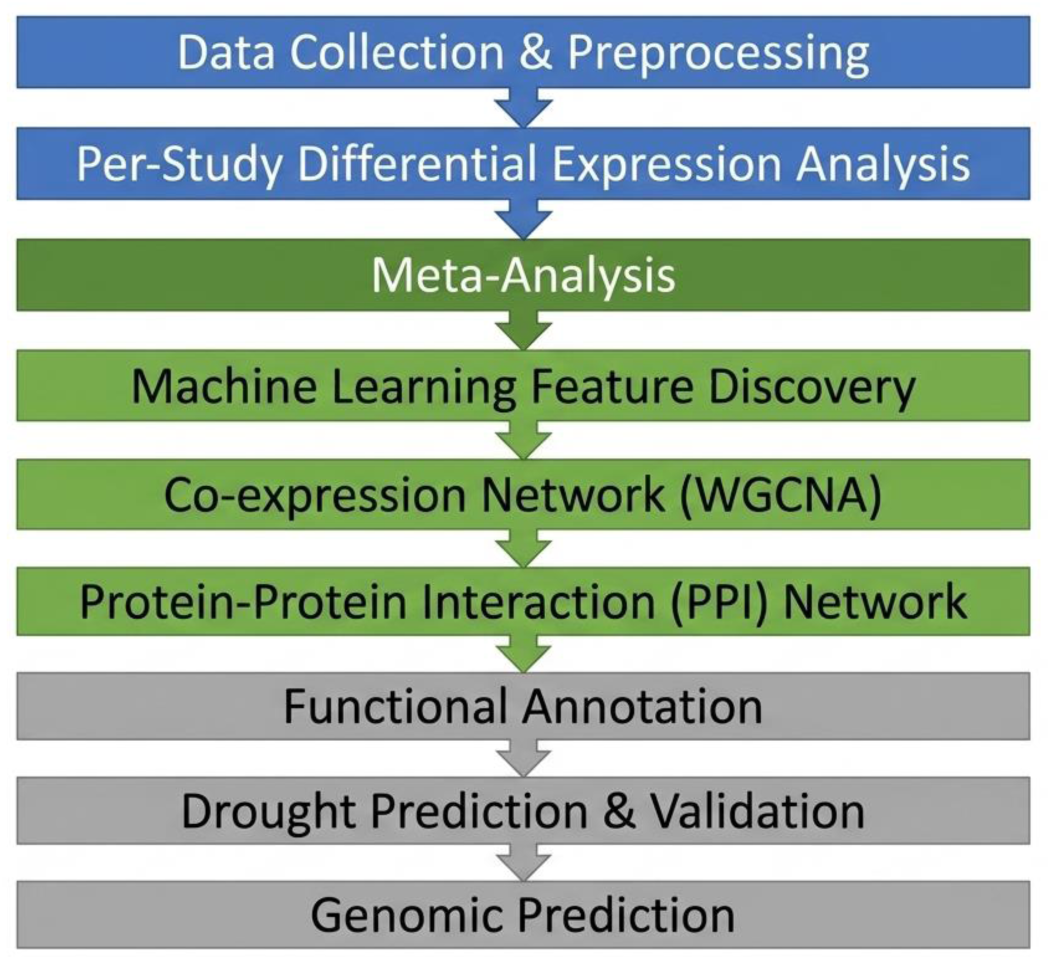
Overview of the analytical pipeline for common bean (*Phaseolus vulgaris*) RNA-seq meta-analysis. Workflow for the analysis integrating per-study differential gene expression analysis, meta-analysis (Fisher’s method and REML), machine learning, WGCNA co-expression network analysis, protein-protein interaction (PPI) network analysis, functional annotation, external drought validation, and genomic prediction validation in cowpea and common bean.

**Figure S2.**
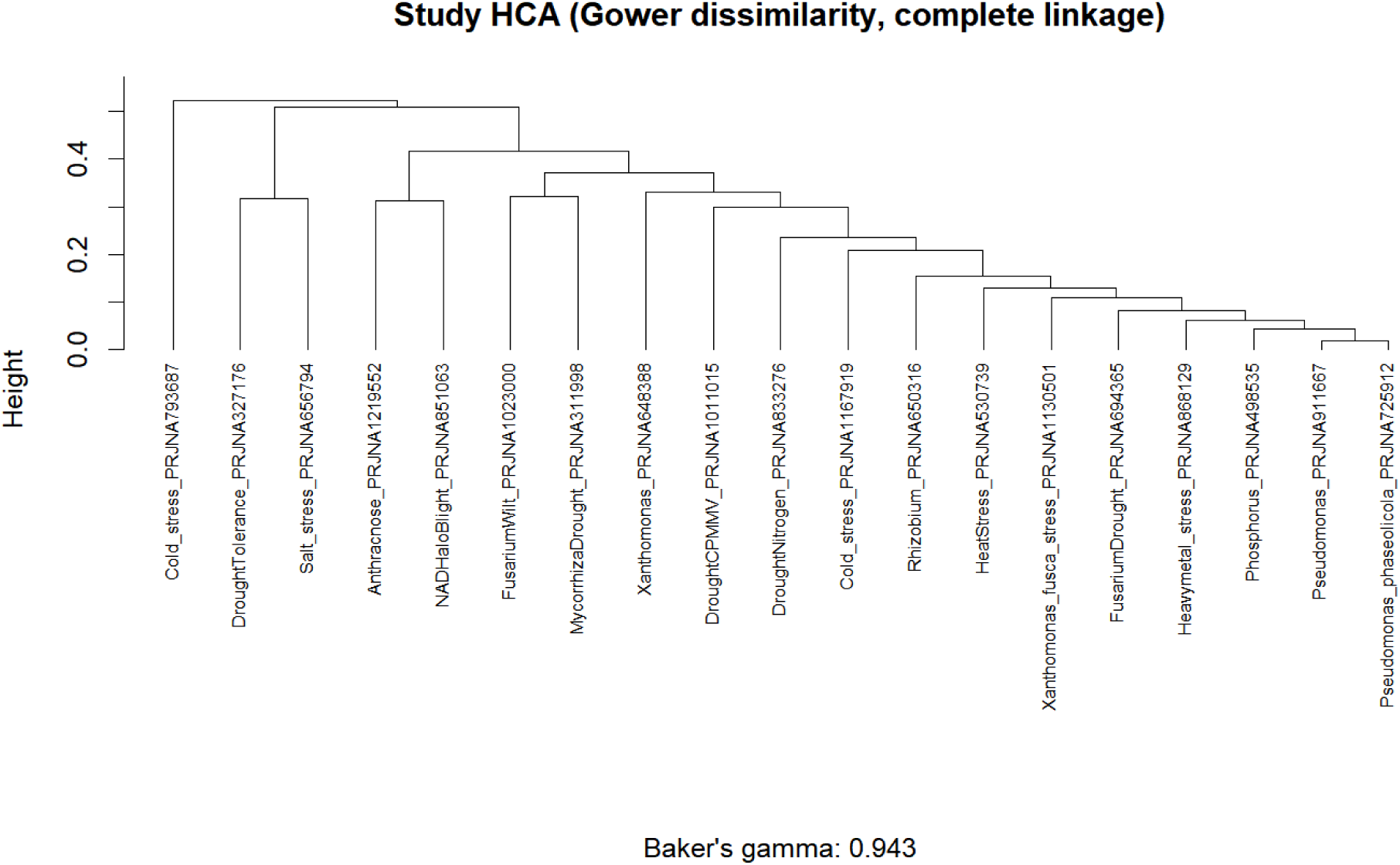
Hierarchical clustering of the 19 included RNA-seq studies based on Gower dissimilarity of binary gene-level differential expression profiles (DEG presence or absence per study), using complete linkage. Baker’s gamma (0.943) reflects high concordance between this complete linkage dendrogram and an alternative dendrogram, indicating the clustering pattern is robust to linkage method choice. Studies group primarily by overall transcriptional scale (total DEG count) rather than by shared category or tissue.

**Figure S3.**
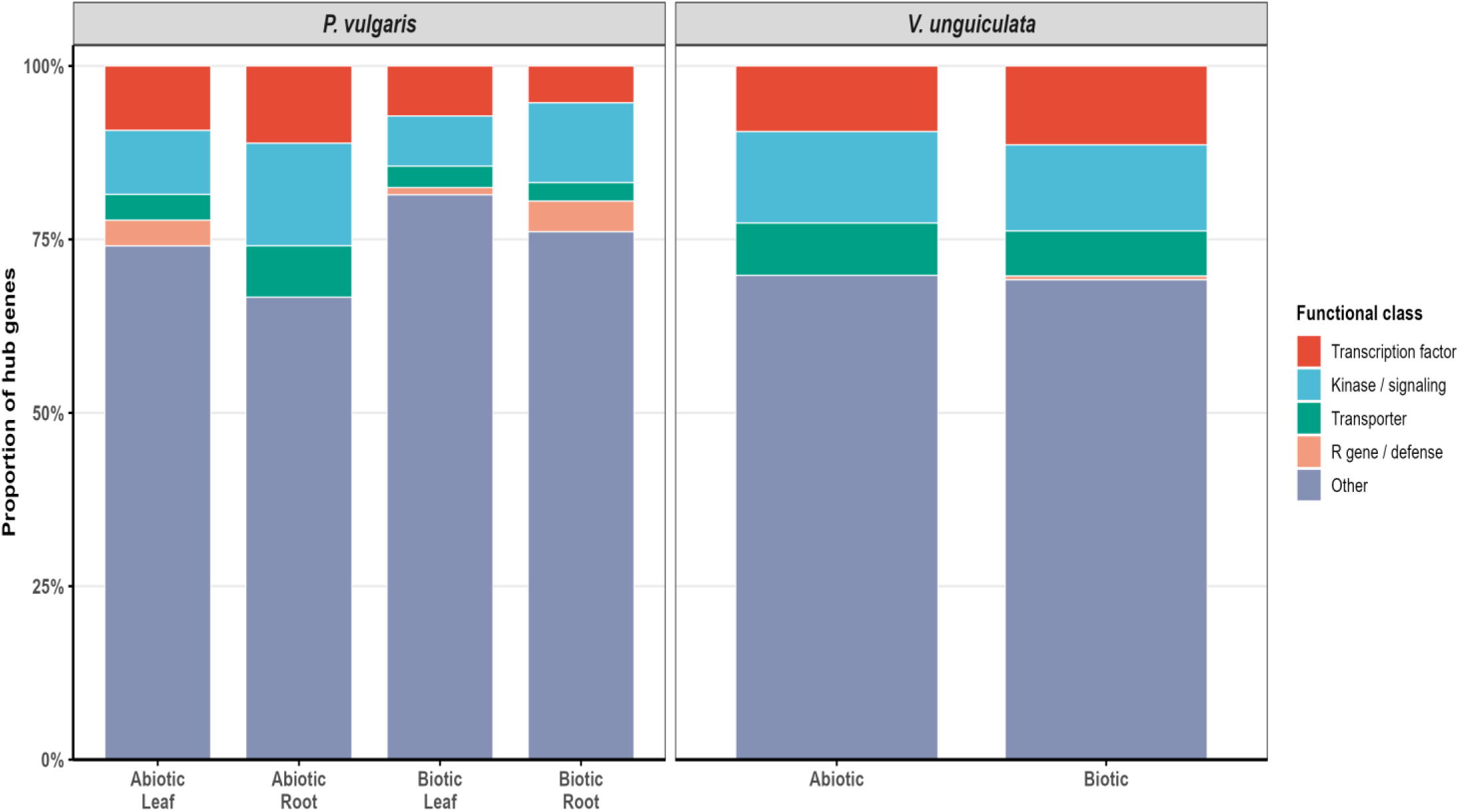
Functional class distribution of ML hub gene orthologs between common bean (*Phaseolus vulgaris*) and cowpea (*Vigna unguiculata*). Proportional distribution of functional classes among ML hub genes in *P. vulgaris* and their confirmed *V. unguiculata* orthologs, split by stress-tissue group. Five functional classes shown: Transcription Factor, Kinase/Signaling, Transporter, R gene/Defense, and Other/Unknown. Only hub genes with confirmed one-to-one cowpea orthologs are included (n = 238; 81.8% of 291 ML hub genes). Functional class assignments are based on Pfam domain annotations and Arabidopsis best-hit descriptions from the *P. vulgaris* v2.1 and *V. unguiculata* v1.2 Phytozome annotation files.

**Figure S4.**
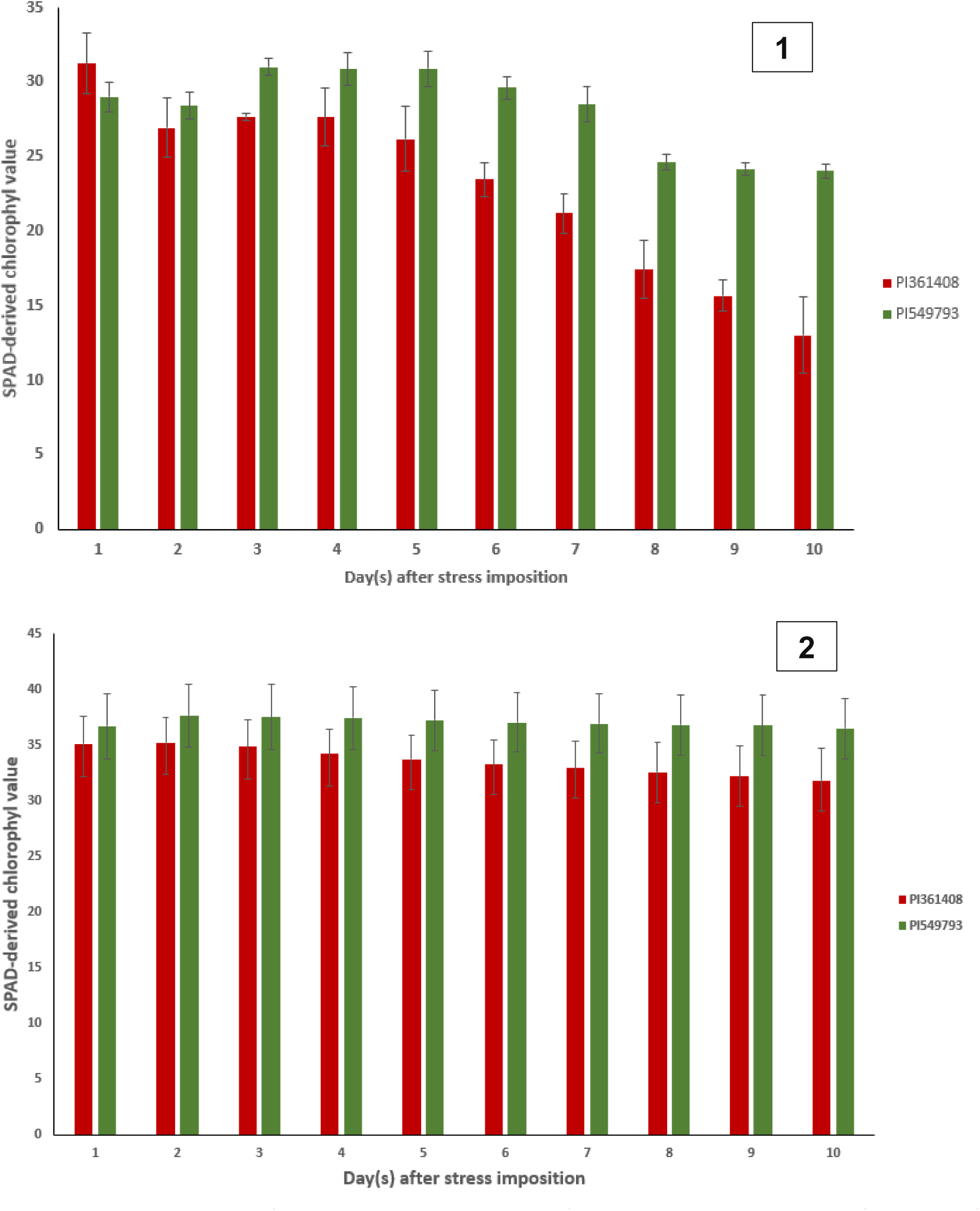
SPAD-derived chlorophyll values over a 10-day period in two *P. vulgaris* genotypes under water-deficit condition in two independent trials (1 and 2). The values for the first trial are from two replicates and the values for the second trial are from four replicates

**Figure S5.**
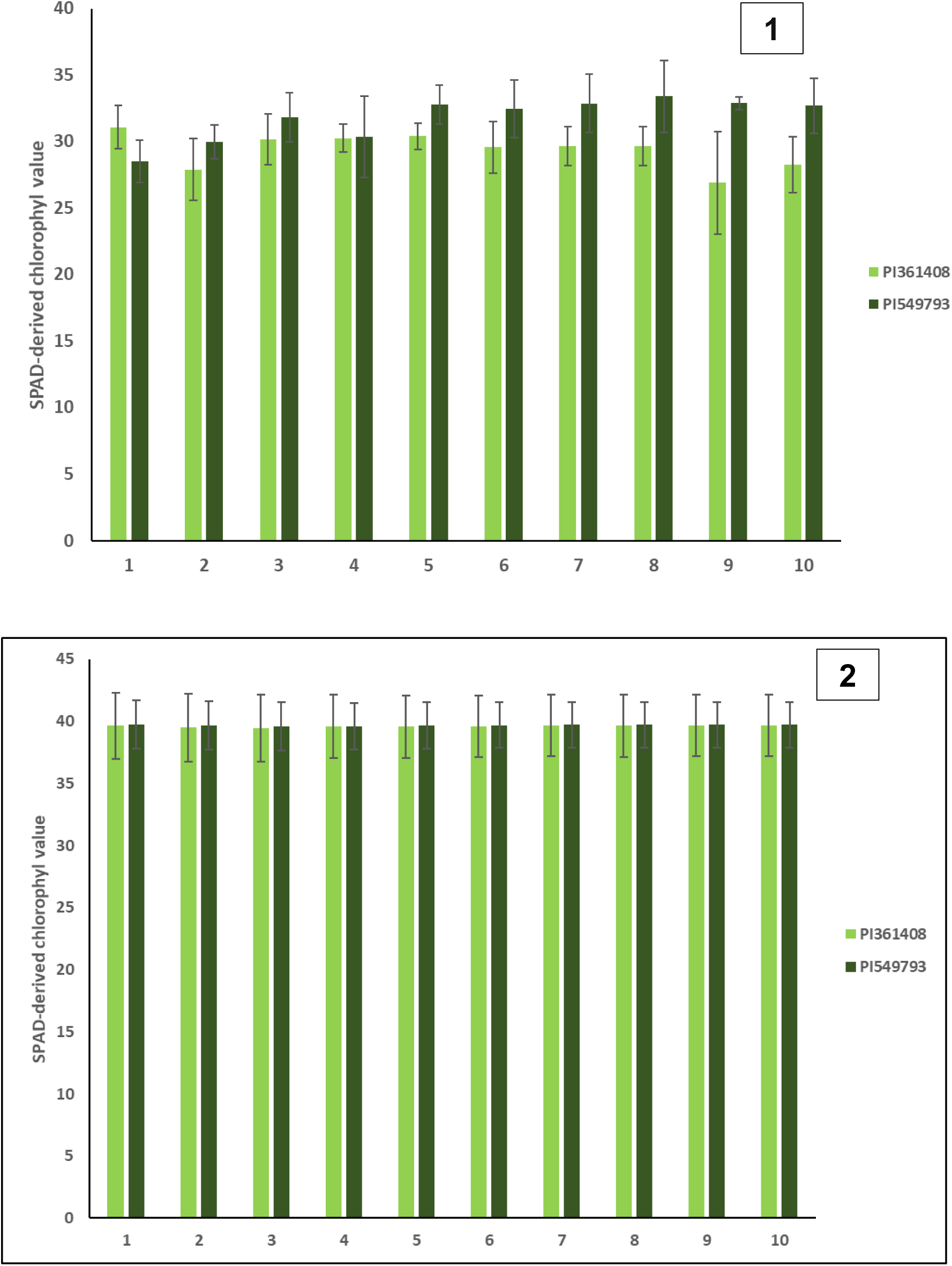
SPAD-derived chlorophyll values over a 10-day period in two *P. vulgaris* genotypes under normal water condition in two independent trials (1 and 2). The values for the first trial are from two replicates and the values for the second trial are from four replicates.

**Figure S6.**
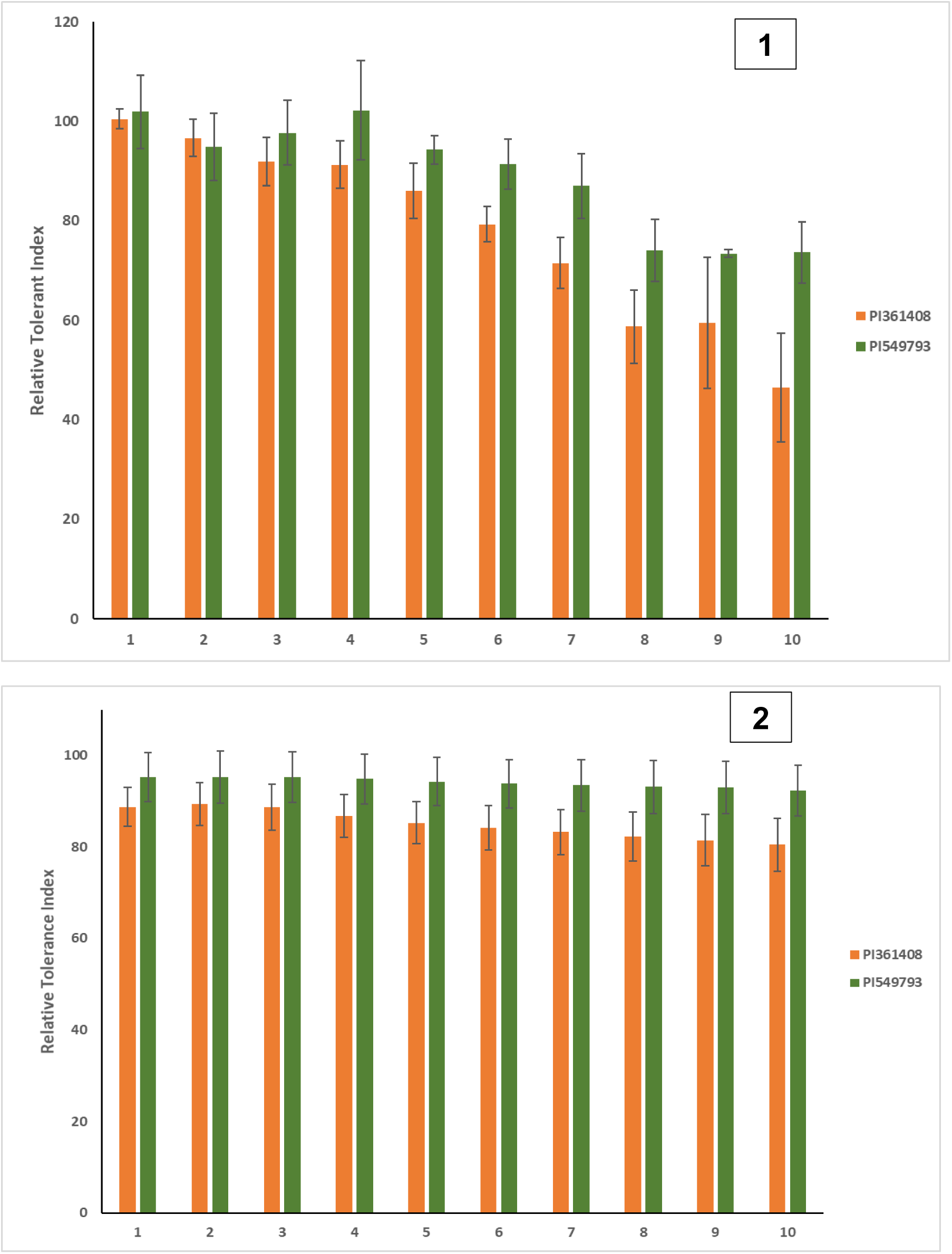
Relative Tolerance Index (RTI) over a 10-day period in two *P. vulgaris* genotypes in two independent trials (1 and 2). Relative tolerance index was obtained by dividing the chlorophyll values from drought-treatment by normal water treatment values. The value was then multiplied by 100 as thus: ((Stress/Non-Stress)*100).

**Figure S7.**
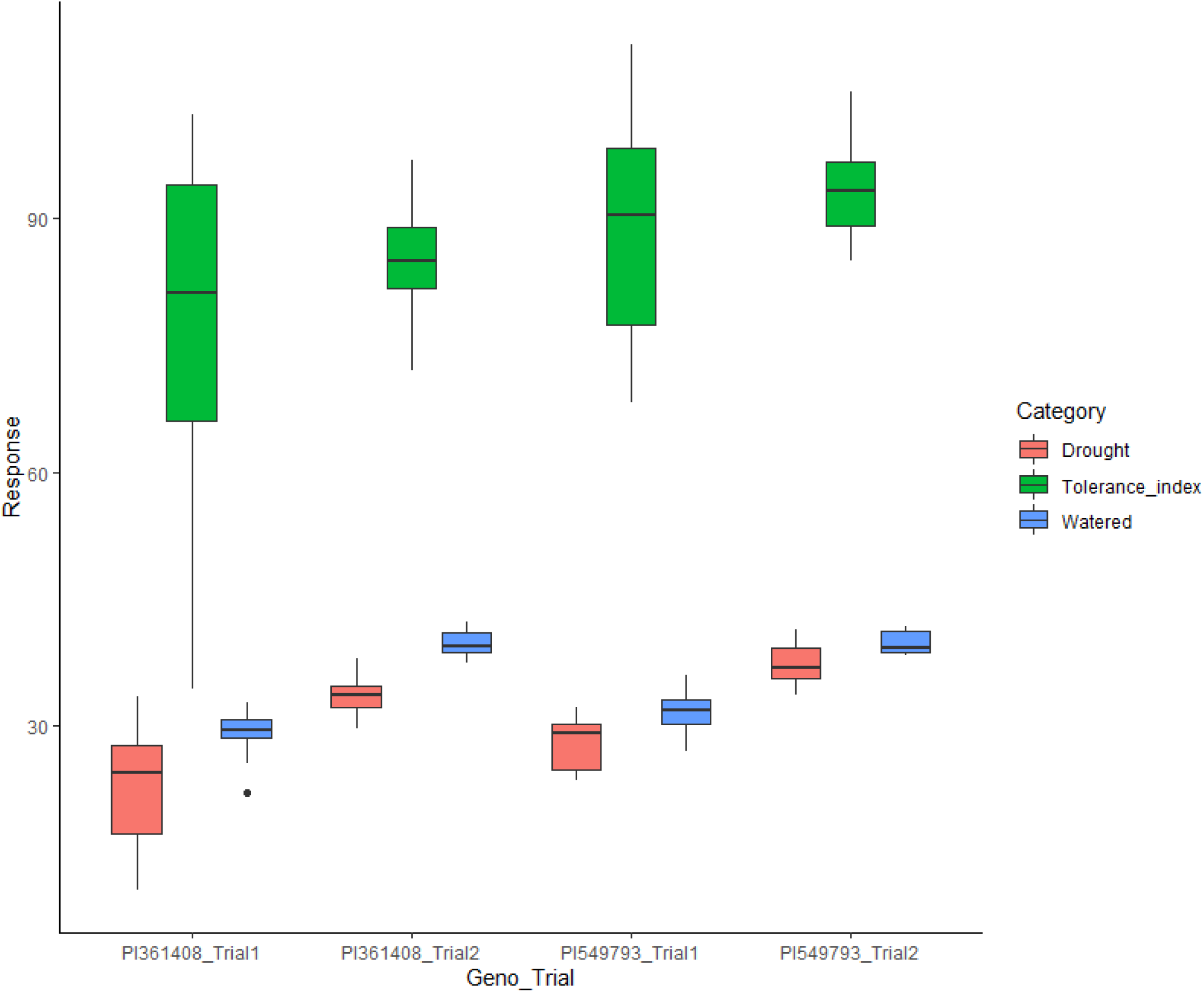
The distribution of SPAD-measured or derived response (chlorophyll content values or tolerance indices) of two *P. vulgaris* genotypes under drought treatment conditions in two independent trials.

**Figure S8.**
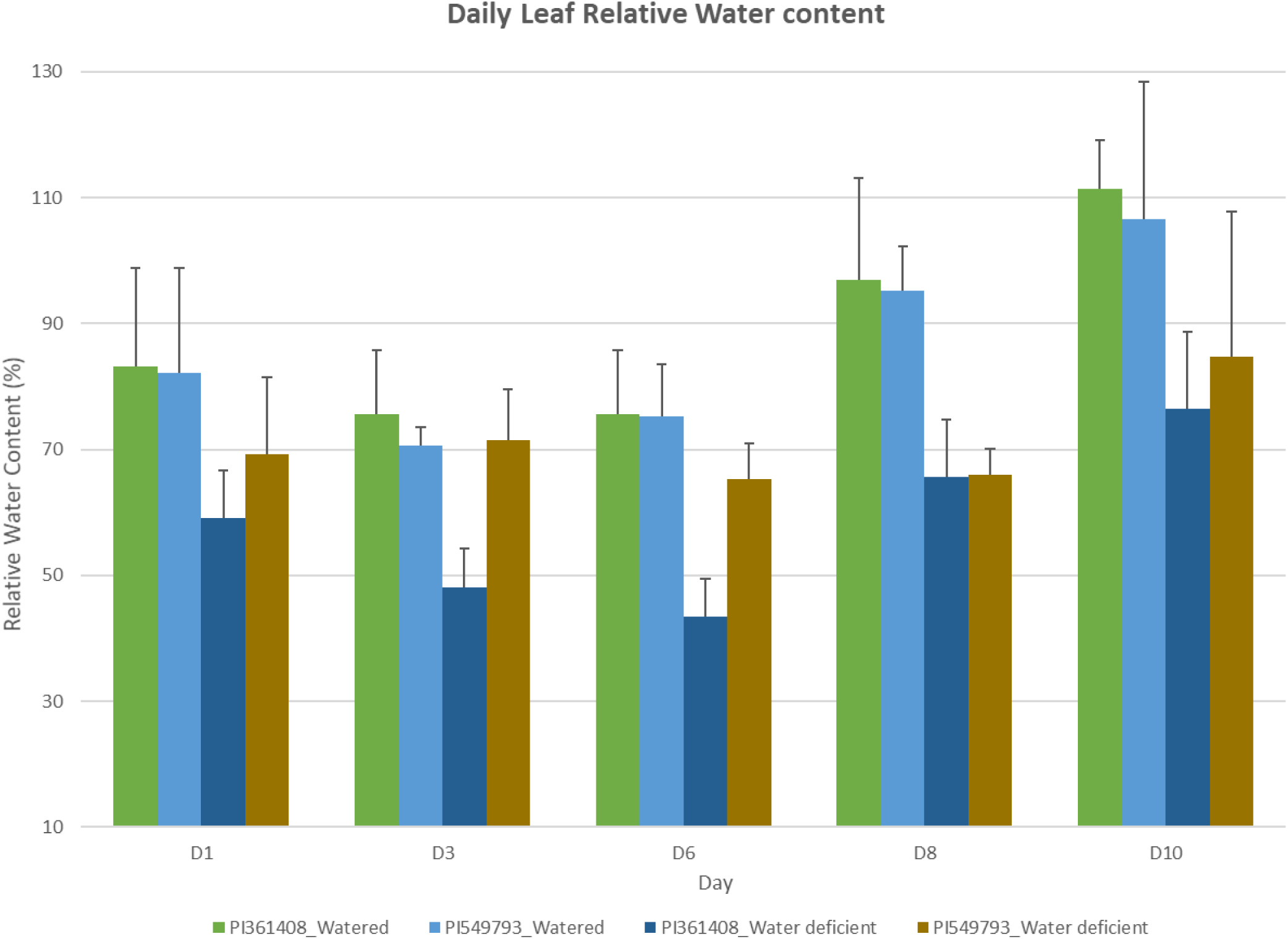
Relative water content values derived over a drought treatment period in two common bean genotypes.

**Figure S9.**
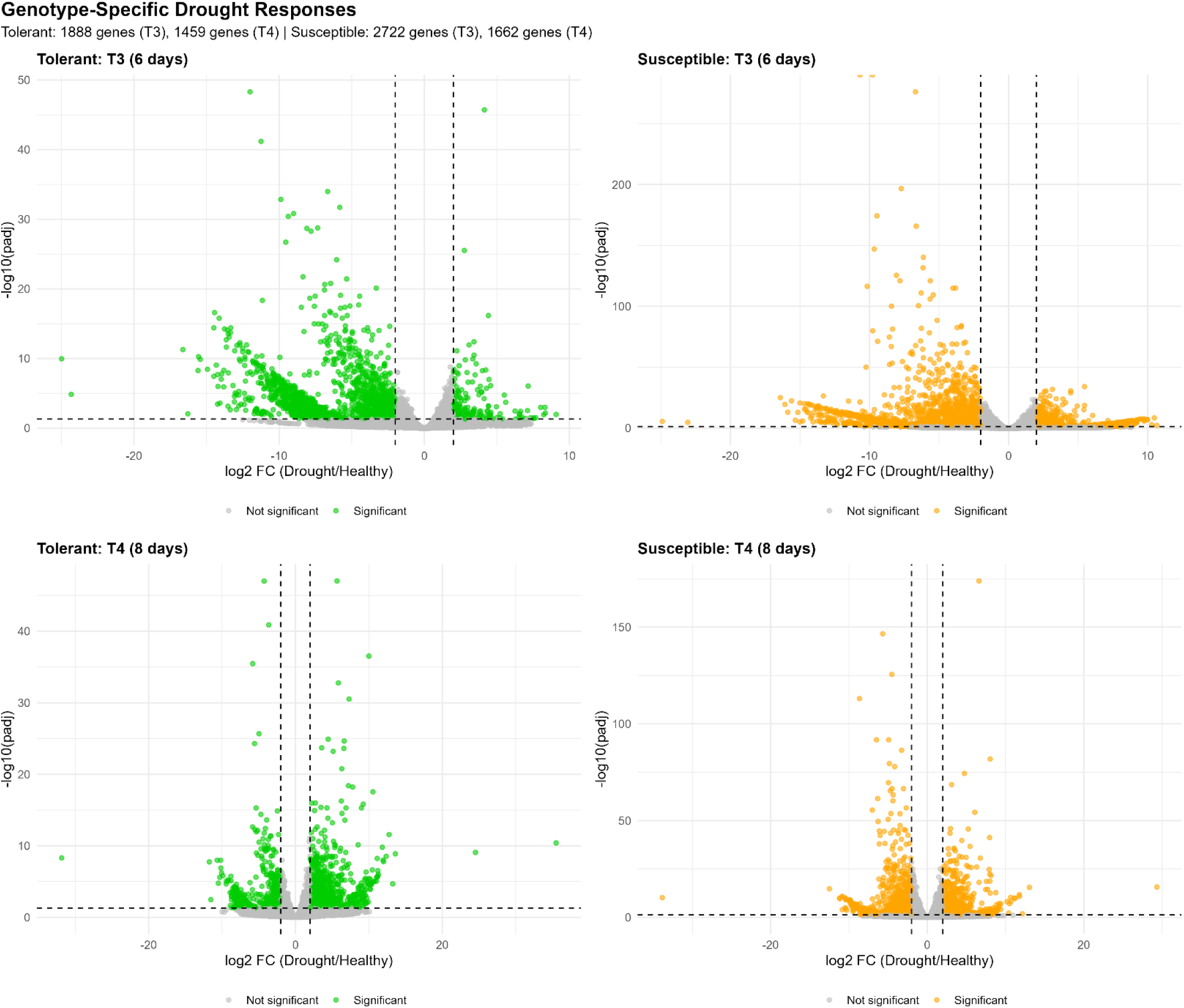
Genotype-specific drought-response for the independent drought experiment. Differential expression (drought vs. healthy) was tested independently within each genotype at each timepoint. A = Tolerant at T3, B = susceptible at T3, C = Tolerant at T4, D = Susceptible at T4. Colored points = significant genes (padj < .05, |log2FC| ≥2); grey points = not significant

## References

Akibode CS, Maredia M (2012) Global and regional trends in production, trade and consumption of food legume crops. Staff Paper 2012–10. Department of Agricultural, Food and Resource Economics, Michigan State University

Al-Azani, S., Alkhnbashi, O. S., Ramadan, E., & Alfarraj, M. (2024). Gene Expression-Based Cancer Classification for Handling the Class Imbalance Problem and Curse of Dimensionality. International journal of molecular sciences, 25(4), 2102. 10.3390/ijms25042102

Aroca, R., Porcel, R., & Ruiz-Lozano, J. M. (2012). Regulation of root water uptake under abiotic stress conditions. Journal of experimental botany, 63(1), 43–57. 10.1093/jxb/err266

Azad M, Tohidfar M, Ghanbari Moheb Seraj R, Mehralian M, Esmaeilzadeh-Salestani K (2024) Identification of responsive genes to multiple abiotic stresses in rice (Oryza sativa): a meta-analysis of transcriptomics data. Sci Rep 14:5463. doi:10.1038/s41598-024-54623-7

Bach-Pages, M., Menon, A., Wadley, B., Castello, A., & Preston, G. M. (2025). RNA-binding proteins orchestrating immunity in plants. The Plant journal : for cell and molecular biology, 123(5), e70433. 10.1111/tpj.70433

Balmer, A., Pastor, V., Gamir, J., Flors, V., & Mauch-Mani, B. (2015). The ’prime-ome’: towards a holistic approach to priming. Trends in plant science, 20(7), 443–452. 10.1016/j.tplants.2015.04.002

Beebe, S.E., I.M. Rao and M.W. Blair, and J.A. Acosta-Gallegos. 2013. Phenotyping common beans for adaptation to drought. Front. Physiol. 4:35. doi:10.3389/fphys.2013.00035

Câmara, C.R., Urrea, C.A., and Schlegel, V.. 2013. Pinto beans (Phaseolus vulgaris L.) as a functional food: Implications on human health. Agriculture 3:90–111.

Chai, H. H., Ho, W. K., Graham, N., May, S., Massawe, F., & Mayes, S. (2017). A Cross-Species Gene Expression Marker-Based Genetic Map and QTL Analysis in Bambara Groundnut. Genes, 8(2), 84. 10.3390/genes8020084

Chowdhury RH, Eti FS, Ahmed R, Das Gupta S, Jhan PK, Islam T, Bhuiyan MAR, Rubel MH, Khayer A (2023) Drought-responsive genes in tomato: meta-analysis of gene expression using machine learning. Sci Rep 13:19374. doi:10.1038/s41598-023-45942-2

Clúa J, Rivero CH, Roda C, Giorgis C, Donna S, Zanetti ME, Blanco FA (2022) Transcriptomic analysis of Mesoamerican and Andean Phaseolus vulgaris accessions revealed mRNAs and lncRNAs associated with strain selectivity during symbiosis. Sci Rep 12:2614. doi:10.1038/s41598-022-06566-0

Conway, J. R., Lex, A., & Gehlenborg, N. (2017). UpSetR: an R package for the visualization of intersecting sets and their properties. *Bioinformatics (Oxford*, England), 33(18), 2938–2940. 10.1093/bioinformatics/btx364

Covarrubias-Pazaran G. (2016). Genome-Assisted Prediction of Quantitative Traits Using the R Package sommer. PloS one, 11(6), e0156744. 10.1371/journal.pone.0156744

da Silva DA, Tsai SM, Chiorato AF, Andrade SCS, Esteves JAF, Recchia GH, Carbonell SAM (2019) Analysis of the common bean (Phaseolus vulgaris L.) transcriptome regarding efficiency of phosphorus use. PLOS ONE 14:e0210428. doi:10.1371/journal.pone.0210428

Dambroz CMS, Aono AH, Costa LC, Novaes E, Pereira WA (2025) Comparative transcriptome analysis provides new insights into the response of common bean to infection by race 65 of Colletotrichum lindemuthianum. PLOS ONE 20:e0314188. doi:10.1371/journal.pone.0314188

Darkwa, K., Ambachew, D., Mohammed, H., Asfaw, A. & Blair, M. W. Evaluation of common bean (Phaseolus vulgaris L.) genotypes for drought stress adaptation in Ethiopia. Crop J. 4, 367–376 (2016).

Diaz, L. M. et al. QTL analyses for tolerance to abiotic stresses in a common bean (Phaseolus vulgaris L.) population. PLoS ONE 13, e0202342 (2018).

Durinck, S., Spellman, P. T., Birney, E., & Huber, W. (2009). Mapping identifiers for the integration of genomic datasets with the R/Bioconductor package biomaRt. Nature protocols, 4(8), 1184–1191. 10.1038/nprot.2009.97

Edwards, S. M., Sørensen, I. F., Sarup, P., Mackay, T. F., & Sørensen, P. (2016). Genomic Prediction for Quantitative Traits Is Improved by Mapping Variants to Gene Ontology Categories in Drosophila melanogaster. Genetics, 203(4), 1871–1883. 10.1534/genetics.116.187161

Fang P, Hu Y, Xia W, Wu X, Sun T, Pandey AK, Ning K, Zhu C, Xu P (2023) Transcriptome dynamics of common bean roots exposed to various heavy metals reveal valuable target genes and promoters for genetic engineering. J Agric Food Chem 71:223–233. doi:10.1021/acs.jafc.2c06301

Foucher J, Ruh M, Préveaux A, Carrère S, Pelletier S, Briand M, Serre RF, Jacques MA, Chen NWG (2020) Common bean resistance to Xanthomonas is associated with upregulation of the salicylic acid pathway and downregulation of photosynthesis. BMC Genomics 21:566. doi:10.1186/s12864-020-06972-6

Friedman, J. H., Hastie, T., & Tibshirani, R. (2010). Regularization Paths for Generalized Linear Models via Coordinate Descent. Journal of Statistical Software, 33(1), 1–22. 10.18637/jss.v033.i01

Galili T. (2015). dendextend: an R package for visualizing, adjusting and comparing trees of hierarchical clustering. *Bioinformatics (Oxford*, England), 31(22), 3718–3720. 10.1093/bioinformatics/btv428

Gihaut C, Brin C, Briand M, Verdier J, Barret M, Roitsch T, Boureau T (2024) Transcriptomic dataset of Phaseolus vulgaris leaves in response to the inoculation of pathogenic Xanthomonas citri pv. fuscans and its type III secretion system-defective mutant hrcV. Data Brief 57:110938.doi:10.1016/j.dib.2024.110938

Gourion, B., Berrabah, F., Ratet, P., & Stacey, G. (2015). Rhizobium-legume symbioses: the crucial role of plant immunity. Trends in plant science, 20(3), 186–194. 10.1016/j.tplants.2014.11.008

Gower, John C. (1971) A General Coefficient of Similarity and Some of Its Properties. Biometrics, 27 (4). pp. 857–871.

Gregorio Jorge, J., Villalobos-López, M. A., Chavarría-Alvarado, K. L., Ríos-Meléndez, S., López-Meyer, M., & Arroyo-Becerra, A. (2020). Genome-wide transcriptional changes triggered by water deficit on a drought-tolerant common bean cultivar. BMC Plant Biol. 20(1), 525. 10.1186/s12870-020-02664-1

Gu, Z., Eils, R., & Schlesner, M. (2016). Complex heatmaps reveal patterns and correlations in multidimensional genomic data. *Bioinformatics (Oxford*, England), 32(18), 2847–2849. 10.1093/bioinformatics/btw313

Karatzoglou, A., Smola, A., Hornik, K., & Zeileis, A. (2004). kernlab - An S4 Package for Kernel Methods in R. Journal of Statistical Software, 11(9), 1–20. 10.18637/jss.v011.i09

Kiranmai, K., Lokanadha Rao, G., Pandurangaiah, M., Nareshkumar, A., Amaranatha Reddy, V., Lokesh, U., Venkatesh, B., Anthony Johnson, A. M., & Sudhakar, C. (2018). A Novel WRKY Transcription Factor, MuWRKY3 (Macrotyloma uniflorum Lam. Verdc.) Enhances Drought Stress Tolerance in Transgenic Groundnut (Arachis hypogaea L.) Plants. Frontiers in plant science, 9, 346. 10.3389/fpls.2018.00346

Kolde, R. (2019). Pheatmap: pretty heatmaps. R package version, 1(2), 726.

Kuhn, M. (2008). Building Predictive Models in R Using the caret Package. Journal of Statistical Software, 28(5), 1–26. 10.18637/jss.v028.i05

Langfelder, P., & Horvath, S. (2008). WGCNA: an R package for weighted correlation network analysis. BMC bioinformatics, 9, 559. 10.1186/1471-2105-9-559

Leek, J. T., Johnson, W. E., Parker, H. S., Jaffe, A. E., & Storey, J. D. (2012). The sva package for removing batch effects and other unwanted variation in high-throughput experiments. *Bioinformatics (Oxford*, England), 28(6), 882–883. 10.1093/bioinformatics/bts034

Leitão ST, Santos C, Araújo SS, Rubiales D, Vaz Patto MC (2021) Shared and tailored common bean transcriptomic responses to combined fusarium wilt and water deficit. Hort Res 8:149. doi:10.1038/s41438-021-00583-2

Liaw, A. and Wiener, M. (2002) Classification and Regression by Randomforest. R News, 2, 18–22.

Lonardi, S., Muñoz-Amatriaín, M., Liang, Q., Shu, S., Wanamaker, S. I., Lo, S., Tanskanen, J., Schulman, A. H., Zhu, T., Luo, M. C., Alhakami, H., Ounit, R., Hasan, A. M., Verdier, J., Roberts, P. A., Santos, J. R. P., Ndeve, A., Doležel, J., Vrána, J., Hokin, S. A., … Close, T. J. (2019). The genome of cowpea (Vigna unguiculata [L.] Walp.). The Plant journal, 98(5), 767–782. 10.1111/tpj.14349

López CM, Alseekh S, Torralbo F, Martínez Rivas FJ, Fernie AR, Amil-Ruiz F, Alamillo JM (2023) Transcriptomic and metabolomic analysis reveals that symbiotic nitrogen fixation enhances drought resistance in common bean. J Exp Bot 74:3203–3219. doi:10.1093/jxb/erad083

Love, M. I., Huber, W., & Anders, S. (2014). Moderated estimation of fold change and dispersion for RNA-seq data with DESeq2. Genome biology, 15(12), 550. 10.1186/s13059-014-0550-8

Madirov, A., Iksat, N., Turarbekova, Z., Abzhalelov, B., & Masalimov, Z. (2025). Competition for Chaperones: A Trade-Off Between Thermotolerance and Antiviral Immunity in Plants. Current issues in molecular biology, 47(11), 957. 10.3390/cimb47110957

Mohi-Ud-Din, M., Talukder, D., Rohman, M., Ahmed, J. U., Jagadish, S. V. K., Islam, T., & Hasanuzzaman, M. (2021). Exogenous Application of Methyl Jasmonate and Salicylic Acid Mitigates Drought-Induced Oxidative Damages in French Bean (Phaseolus vulgaris L.). Plants (Basel, Switzerland), 10(10), 2066. 10.3390/plants10102066

Muñoz-Perea, C.G., Teran, H., Allen, R.G., Wright, J.L., Westermann, D.T., and Singh, S.P.. 2006. Selection for drought resistance in dry bean landraces and cultivars. Crop Sci. 46:2111–2120. doi:10.2135/cropsci2006.01.0029

Niron H, Barlas N, Salih B, Türet M (2020) Comparative transcriptome, metabolome, and ionome analysis of two contrasting common bean genotypes in saline conditions. Front Plant Sci 11:599501. doi:10.3389/fpls.2020.599501

Oblessuc PR, Bridges DF, Melotto M (2022) Pseudomonas phaseolicola preferentially modulates genes encoding leucine-rich repeat and malectin domains in the bean landrace G2333. Planta 256:25. doi:10.1007/s00425-022-03943-x

Panahi B (2024) Transcriptome signature for multiple biotic and abiotic stress in barley (Hordeum vulgare L.) identifies using machine learning approach. Curr Plant Biol 40:100416. doi:10.1016/j.cpb.2024.100416

Pandey, E., Kumari, R., Faizan, S., & Pandey, S. (2025). Linking the interaction of Salicylates and Jasmonates for stress resilience in plants. Stress biology, 5(1), 64. 10.1007/s44154-025-00250-9

Pandey, S., Fartyal, D., Agarwal, A., Shukla, T., James, D., Kaul, T., Negi, Y. K., Arora, S., & Reddy, M. K. (2017). Abiotic Stress Tolerance in Plants: Myriad Roles of Ascorbate Peroxidase. Frontiers in plant science, 8, 581. 10.3389/fpls.2017.00581

Patro, R., Duggal, G., Love, M. I., Irizarry, R. A., & Kingsford, C. (2017). Salmon provides fast and bias-aware quantification of transcript expression. Nature methods, 14(4), 417–419. 10.1038/nmeth.4197

Pereira, W. J., Melo, A., Coelho, A., Rodrigues, F. A., Mamidi, S., Alencar, S. A., Lanna, A. C., Valdisser, P., Brondani, C., Nascimento-Júnior, I., Borba, T., & Vianello, R. P. (2020). Genome-wide analysis of the transcriptional response to drought stress in root and leaf of common bean. Genet Mol Biol. 43(1), e20180259.

Pieterse, C. M., Leon-Reyes, A., Van der Ent, S., & Van Wees, S. C. (2009). Networking by small-molecule hormones in plant immunity. Nature chemical biology, 5(5), 308–316. 10.1038/nchembio.164

Polania, J., Poschenrieder, C., Rao, I., & Beebe, S. (2016). Estimation of phenotypic variability in symbiotic nitrogen fixation ability of common bean under drought stress using 15N natural abundance in grain. Eur. J. Agron. 79, 66–73.

Rao, I. M., Beebe, S., Polania, J., Grajales, M. A. & García, R. Differences in drought resistance of advanced lines developed for the last 3 decades. In Annual Report 2006. Project IP-1: Bean Improvement for the Tropics. (CIAT, Cali, Colombia, 2006). pp 2–6.

Ravelombola W, Dong L, Barickman TC, Xiong H, Manley A, Cason J, Pham H, Zia B, Mou B, Shi A (2023) Genetic architecture of salt tolerance in cowpea (Vigna unguiculata (L.) Walp.) at seedling stage using a whole genome resequencing approach. Int J Mol Sci 24:15281. doi:10.3390/ijms242015281

Ravelombola W, Xiong H, Bhattarai G, Manley A, Cason J, Pham H, Zia B, Mou B, Shi A (2025) Genome-wide association study for drought tolerance in cowpea (Vigna unguiculata (L.) Walp.) at seedling stage using a whole genome resequencing approach. Int J Mol Sci 26:5478. doi:10.3390/ijms26125478

Recchia GH, Konzen ER, Cassieri F, Caldas DGG, Tsai SM (2018) Arbuscular mycorrhizal symbiosis leads to differential regulation of drought-responsive genes in tissue-specific root cells of common bean. Front Microbiol 9:1339. doi:10.3389/fmicb.2018.01339

Robin, X., Turck, N., Hainard, A., Tiberti, N., Lisacek, F., Sanchez, J. C., & Müller, M. (2011). pROC: an open-source package for R and S+ to analyze and compare ROC curves. BMC bioinformatics, 12, 77. 10.1186/1471-2105-12-77

Rosales-Serna, R., Kohashi-Shibata, J., Acosta-Gallegos, J.A., Trejo-Lopez, C., Ortiz-Cereceres, J., and Kelly, J.D.. 2004. Biomass distribution, maturity acceleration, and yield in drought-stressed common bean cultivars. Field Crops Res. 85:203–211. doi: 10.1016/S0378-4290(03)00161-8

Sanchez-Munoz R, Depaepe T, Samalova M, Hejatko J, Zaplana I, Van Der Straeten D (2025) Machine-learning meta-analysis reveals ethylene as a central component of the molecular core in abiotic stress responses in Arabidopsis. Nat Commun 16:4778. doi:10.1038/s41467-025-59542-3

Schmutz, J., McClean, P. E., Mamidi, S., Wu, G. A., Cannon, S. B., Grimwood, J., Jenkins, J., Shu, S., Song, Q., Chavarro, C., Torres-Torres, M., Geffroy, V., Moghaddam, S. M., Gao, D., Abernathy, B., Barry, K., Blair, M., Brick, M. A., Chovatia, M., Gepts, P., … Jackson, S. A. (2014). A reference genome for common bean and genome-wide analysis of dual domestications. Nat. Genet. 46, 707–713

Shi A, Xiong H, Michaels TE, Chen S (2025) Genome and GWAS analyses for soybean cyst nematode resistance in USDA world-wide common bean (Phaseolus vulgaris) germplasm. Front Plant Sci 16:1520087. doi:10.3389/fpls.2025.1520087

Shintani M, Bono H (2025) Meta-analysis of public RNA-sequencing data of drought and salt stresses in different phenotypes of resistant and susceptible Oryza sativa cultivars. Quantitative Plant Biol 6:e27. doi:10.1017/qpb.2025.10020

Smith, M. R., Veneklaas, E., Polania, J., Rao, I. M., Beebe, S. E., & Merchant, A. 2019. Field drought conditions impact yield but not nutritional quality of the seed in common bean (Phaseolus vulgaris L.). PloS one, 14(6), e0217099.

Soltani A, Weraduwage SM, Sharkey TD, Lowry DB (2019) Elevated temperatures cause loss of seed set in common bean (Phaseolus vulgaris L.) potentially through the disruption of source-sink relationships. BMC Genomics 20:312. doi:10.1186/s12864-019-5669-2

Soneson, C., Love, M. I., & Robinson, M. D. (2015). Differential analyses for RNA-seq: transcript-level estimates improve gene-level inferences. F1000R*esearch*, *4*, 1521. 10.12688/f1000research.7563.2

Szklarczyk, D., Kirsch, R., Koutrouli, M., Nastou, K., Mehryary, F., Hachilif, R., Gable, A. L., Fang, T., Doncheva, N. T., Pyysalo, S., Bork, P., Jensen, L. J., & von Mering, C. (2023). The STRING database in 2023: protein-protein association networks and functional enrichment analyses for any sequenced genome of interest. Nucleic acids research, 51(D1), D638–D646. 10.1093/nar/gkac1000

Tang W, Li Z, Xu Z, Sui X, Liang L, Xiao J, Song X, Sun B, Huang Z, Lai Y, Wang C, Tang Y, Li H (2025) Transcriptomic and metabolomic analysis reveal the cold tolerance mechanism of common beans under cold stress. BMC Plant Biol 25:340. doi:10.1186/s12870-025-06333-z

Tarun, J. A., Mauleon, R., Arbelaez, J. D., Catausan, S., Dixit, S., Kumar, A., Brown, P., Kohli, A., & Kretzschmar, T. (2020). Comparative Transcriptomics and Co-Expression Networks Reveal Tissue- and Genotype-Specific Responses of qDTYs to Reproductive-Stage Drought Stress in Rice (Oryza sativa L.). Genes, 11(10), 1124. 10.3390/genes11101124

Torres, G. A., Pflieger, S., Corre-Menguy, F., Mazubert, C., Hartmann, C., & Lelandais-Brière, C. 2006. Identification of novel drought-related mRNAs in common bean roots by differential display RT-PCR. Plant science, 171:300–307.

VanRaden PM (2008) Efficient methods to compute genomic predictions. J Dairy Sci 91:4414–4423. doi:10.3168/jds.2007-0980

Verma, V., Ravindran, P., & Kumar, P. P. (2016). Plant hormone-mediated regulation of stress responses. BMC Plant Biol. 16, 86. 10.1186/s12870-016-0771-y

Viechtbauer, W. (2010). Conducting Meta-Analyses in R with the metafor Package. Journal of Statistical Software, 36(3), 1–48. 10.18637/jss.v036.i03

Villordo-Pineda, E., González-Chavira, M., Giraldo-Carbajo, P., Acosta-Gallegos, J. & Caballero-Pérez, J. Identification of novel drought-tolerant-associated SNPs in common bean (Phaseolus vulgaris). Front. Plant Sci. 6, 546 (2015).

Wright, M. N., & Ziegler, A. (2017). ranger: A Fast Implementation of Random Forests for High Dimensional Data in C++ and R. Journal of Statistical Software, 77(1), 1–17. 10.18637/jss.v077.i01

Wu J, Wang L, Li L, Wang S. 2014. De Novo Assembly of the Common Bean Transcriptome Using Short Reads for the Discovery of Drought-Responsive Genes. PLoS One 9(10): e109262

Wu X, Chen S, Zhang Z, Zhou W, Sun T, Ning K, Xu M, Ke X, Xu P (2024) A viral small interfering RNA-host plant mRNA pathway modulates virus-induced drought tolerance by enhancing autophagy. Plant Cell 36:3219–3236. doi:10.1093/plcell/koae158

Wu, J., Chen, J., Wang, L., & Wang, S. 2017. Genome-Wide Investigation of WRKY Transcription Factors Involved in Terminal Drought Stress Response in Common Bean. Front. Plant Sci. 8, 380. 10.3389/fpls.2017.00380

Wu, T., Hu, E., Xu, S., Chen, M., Guo, P., Dai, Z., Feng, T., Zhou, L., Tang, W., Zhan, L., Fu, X., Liu, S., Bo, X., & Yu, G. (2021). clusterProfiler 4.0: A universal enrichment tool for interpreting omics data. Innovation (Cambridge (Mass.)), 2(3), 100141. 10.1016/j.xinn.2021.100141

Yang X, Liu C, Li M, Li Y, Yan Z, Feng G, Liu D (2023) Integrated transcriptomics and metabolomics analysis reveals key regulatory network that response to cold stress in common bean (Phaseolus vulgaris L.). BMC Plant Biol 23:85. doi:10.1186/s12870-023-04094-1

Yang, M., Gao, A., Chan, T. Y., Rehman, H. M., Nawaz, S., Bashir, S., Mahmood, N., Ullah, M. W., Long, C., Lam, H. M., & Arif, M. (2026). Molecular Chaperone Networks in Plants: Maintaining Proteostasis and Enhancing Stress Resilience for Crop Improvement. Plant, cell & environment, 10.1111/pce.70629. Advance online publication. https://doi.org/10.1111/pce.70629

Yang, Y., Cui, Q., Qin, J., Ravelombola, W., Bhattarai, G., Zia, B., Samuel, O. D., Imamura, S., Archbold, J. S., & Shi, A. (2019) Evaluation of Drought Tolerance in Common Bean at Seedling Stage [Abstract]. ASA, CSSA and SSSA International Annual Meetings (2019), San Antonio, TX. https://scisoc.confex.com/scisoc/2019am/meetingapp.cgi/Paper/122167

Ye, S., Li, J., & Zhang, Z. (2020). Multi-omics-data-assisted genomic feature markers preselection improves the accuracy of genomic prediction. Journal of animal science and biotechnology, 11(1), 109. 10.1186/s40104-020-00515-5

Zia B, Shi A, Olaoye D, Xiong H, Ravelombola W, Gepts P, Schwartz HF, Brick MA, Otto K, Ogg B, Chen S (2022) Genome-wide association study and genomic prediction for bacterial wilt resistance in common bean (Phaseolus vulgaris) core collection. Front Genet 13:853114. doi:10.3389/fgene.2022.853114

